# Discrete Subdomains Establish Epigenetic Diversity in Subtelomeric Heterochromatin

**DOI:** 10.1101/2025.09.25.678047

**Authors:** Agnisrota Mazumder, John Cooper, Can Goksal, Rosario Y. J. Brockhausen, Jonas M. Schauß, Jasbeer S. Khanduja, Richard I. Joh, Ahmed A. A. Amine, Johannes J. Groos, Junko Kanoh, Mo Motamedi, Ilya J. Finkelstein, Bassem Al-Sady, Sigurd Braun

**Author notes:** These authors contributed equally (Can Goksal, Rosario Y. J. Brockhausen, Jonas M. Schauß).

## Abstract

Subtelomeres are imperfect repeats adjacent to telomeres that are transcriptionally repressed by heterochromatin. Although essential for genome integrity, their repetitive nature has thwarted dissection of local heterochromatin assembly and maintenance mechanisms. By engineering *Schizosaccharomyces pombe* strains carrying fluorescent reporters at a single subtelomere, we uncovered distinct subdomains. These subdomains have different silencing requirements: Telomere-proximal regions rely on canonical shelterin- or RNAi-dependent nucleation pathways, whereas telomere-distal regions involve nucleosome remodelers, histone chaperones, and boundary-associated factors. Subdomains also exhibit discrete epigenetic states emerging both at homologous loci on different chromosome arms and along the same subtelomeric sequence. We document these epigenetic states using multi-generational live imaging and targeted perturbations. These analyses show that subtelomeric subdomains display position-specific, clonally variable silencing across a spectrum from robust to fragile epigenetic states. Interestingly, structural variants, common in subtelomeric sequences across eukaryotes, dictate local epigenetic stability. These findings reveal that subtelomeres, long recognized for their sequence variability, form a dynamic mosaic of coexisting epigenetic domains. This implies a wide range of gene-repressive regulatory logic at the chromosome ends, from environmental responsiveness to silencing stability.

---

Gene-repressing heterochromatin is a highly conserved feature marked by the methylation (me) of histone 3 (H3) at lysine 9 (K9). This histone modification is deposited by Suv39-family methyl-transferases, such as Suv39h1 and Suv39h2 in mammalian systems or Clr4 in the fission yeast *S. pombe* (Padeken et al., 2022). Heterochromatin formation is initiated when Suv39 methyltransferases are guided to specific sites by non-coding RNAs or DNA-bound factors. Once nucleated by this mechanism, heterochromatin expands into adjacent regions via a DNA sequence-independent mechanism termed spreading. This expansion depends on conserved positive feedback involving Suv39, its enzymatic product, and reader proteins recognizing H3K9me, including members of the Heterochromatin Protein 1 (HP1) family (Allshire and Madhani 2018; Bell et al. 2023; Grewal 2023; Moazed 2009). Local three-dimensional (3D) contacts between nucleosomes are critical for this positive feedback across systems, facilitating both spreading and maintenance of the heterochromatic domains (Erdel and Greene 2016; Owen et al. 2023). In both yeast and metazoans, such mechanisms typically support only short-range spreading, rarely exceeding 10 kb (Abdulla et al. 2022; Dodd et al. 2007; Erdel and Greene 2016; Hamali et al. 2023). In *S. pombe*, this constraint is evident when heterochromatin nucleates and spreads within a euchromatic region, either through ectopic heterochromatin assembly (Audergon et al. 2015; Greenstein et al. 2018, 2020; Ragunathan et al. 2015) or at native heterochromatin islands (Zofall et al. 2012). However, assessing spreading in constitutive heterochromatin is more complex, as these domains often contain multiple distinct nucleation sites (Hansen et al. 2011; Jia et al. 2004; Khanduja et al. 2024) and are insulated by defined DNA elements in yeast and mammals (CTCF) (Cam et al. 2005). Consequently, it remains unclear whether spreading in these regions is limited or whether it can propagate more extensively than in euchromatic environments.

Such long-range spreading is best assessed in simple model systems with defined and minimally redundant heterochromatin pathways. *S. pombe* has been an outstanding model system for dissecting mechanisms of constitutive heterochromatin formation and addressing these questions. This is due to its limited and spatially defined heterochromatin content making up only 2% of its genome, the low redundancy of *trans* acting factors, and its facile genetics. Much progress has been made in uncovering mechanisms of heterochromatin formation at domains with defined nucleation sites, in particular the MAT locus and the pericentromeres. However, the subtelomeric regions, which present the most conserved and largest heterochromatin domains extending over 50 kb (Cam et al. 2005; Kanoh et al. 2005; Tashiro et al. 2017), have been proven more difficult to investigate. Subtelomeres lie adjacent to telomeres but are structurally distinct, consisting of mosaic-like arrays of common sequence elements that occur in variable copy numbers rather than the short tandem repeats present at telomeric regions. This structural complexity, combined with their repeated occurrence across chromosomes, has hindered efforts to precisely define local heterochromatin spreading dynamics. Although subtelomeres are hotspots of genome evolution, subtelomeric heterochromatin plays a protective role by suppressing excessive recombination and restricting the expression of transcripts whose misregulation has been linked to disease (Kanoh 2023; Yip and Picketts 2003). Despite this important function, the rules governing subtelomeric heterochromatin formation, dynamics, and epigenetic stability remain poorly understood. In *S. pombe*, as in other systems, subtelomeres lack known internal nucleation sites and instead rely on heterochromatin nucleation near chromosome ends, from which H3K9me-marked domains are thought to extend. Although the molecular organization of telomeres varies across model systems, heterochromatin nucleation at chromosomal ends, either within telomere repeats or adjacent subtelomeric regions, appear to be evolutionary conserved across fungi, plants, and animals (Vaquero-Sedas and Vega-Palas 2023; Linardopoulou et al. 2005; Hocher and Taddei 2020).

In *S. pombe*, subtelomeric heterochromatin assembles at the subtelomeric homologous (SH) sequences, which are present on most chromosomal arms. Ranging from 33–55 kb, the SH sequences are composed of imperfect repeats and multiple gene-containing segments that share ∼90% identity between chromosomal arms, with occasional insertions and deletions (Kanoh 2023; Oizumi et al. 2021; Yadav et al. 2021). SH regions lack DNA-encoded boundary elements and instead exhibit gradual transitions into adjacent subtelomeric unique (SU) regions, which are devoid of heterochromatic marks (Tashiro et al. 2017; Yadav et al. 2021). Two distinct nucleation pathways initiate heterochromatin formation near the telomeric repeats through the recruitment of Clr4. The first relies on RNA interference (RNAi) and nucleates at repeats present in *tlh1–4* genes, ancient duplications of RecQ helicase-derived genes (Hansen et al. 2006). These repeats drive RNAi-mediated nucleation at pericentromeres, the MAT locus, and subtelomeres via the RNA-induced transcriptional silencing (RITS) complex, in which the scaffold protein Tas3 that bridges Argonaute (Ago1) and the chromodoman protein Chp1 (Cam et al. 2005; Hansen et al. 2006; Kanoh et al. 2005; Oizumi et al. 2021; Verdel et al. 2004). The second pathway depends on the telomere-protecting shelterin complex, which is targeted to telomere-proximal sequences by Taz1 and recruits Clr4 and the Snf2-like remodeler/HDAC-containing repressor complex (SHREC) via Ccq1 (van Emden et al. 2019; Kanoh et al. 2005; Sugiyama et al. 2007; Miyoshi et al. 2008; Zofall et al. 2016; Wang et al. 2016; Cooper et al. 1997). In addition, the inner nuclear membrane protein Lem2 facilitates SHREC recruitment and acts redundantly with shelterin and RNAi (Barrales et al. 2016; Tange et al. 2016; Banday et al. 2016).

The relatively large size and the absence of known boundary elements raise key questions about how such extended heterochromatin is assembled and maintained. In one model, SH may form a single continuous domain. Here, RNAi- and shelterin-dependent pathways seed heterochromatin at the telomere-proximal end, resulting in its propagation toward the SH/SU boundary. This process potentially requires mechanisms that exclude heterochromatin antagonists such as Epe1 (Ayoub et al. 2003). Alternatively, multiple independent nucleation events may occur along the SH region, generating distinct subdomains. To overcome the challenges of SH sequence similarity within and across chromosome arms (Oizumi et al. 2021), we integrated a series of gene silencing reporters into a previously generated yeast strain, which only contains a single SH domain (Tashiro et al. 2017). This approach enabled the detailed dissection of the molecular characteristics, epigenetic behavior, and genetic requirements governing SH heterochromatin. We find evidence for multiple, epigenetically distinct subdomains within the SH region that initiate heterochromatin formation independently of previously described, canonical molecular mechanisms.

## Results

### The SH domain carries a heterogeneous chromatin landscape

To determine whether the ∼50 kb heterochromatin domain assembled over SH sequences forms a contiguous domain or comprises discrete domains with independent nucleation sites, we analyzed a fission yeast strain in which all but one SH domain were replaced by marker genes (*his7, ura4*) (Tashiro et al. 2017). Specifically, we used the *SD4[2R+*] strain that carries a single SH domain on the right arm of chromosome II, while the remaining SH sequences have been removed (Figure 1a). Heterochromatin dosage overall is not altered in this strain, as SH-independent heterochromatin of comparable size is still formed on the other chromosome ends (Tashiro et al. 2017). Chromatin immunoprecipitation followed by quantitative PCR (ChIP–qPCR) across individual subtelomeric genes revealed strong enrichment of tri- and di-methylated H3K9 (H3K9me3, H3K9me2) within a ∼10 kb region near the telomere (Figure 1b, c). This region includes the RNAi-dependent nucleation site at *tlh2*, which shares sequence similarity with pericentromeric *dg-dh* repeats and the *cenH* element at the mating type locus (Kanoh et al. 2005). We found that H3K9me3 levels peak near the *tlh2* nucleation site, while H3K9me2 reaches maximum levels more distally. Beyond this region, we also observed that H3K9me levels decline sharply but remain detectable, consistent with previous observations (Kanoh et al. 2005; Tashiro et al. 2017). To examine euchromatic features, we measured acetylation of histone H3 at lysine 14 (H3K14ac), a mark associated with active transcription (Wiren 2005). H3K14ac displays an inverse pattern to H3K9me, with higher levels in telomere-distal regions and sporadic enrichment at individual SH genes (Figure 1d). These chromatin profiles suggest a non-uniform landscape within the SH domain.

**Figure 1:**
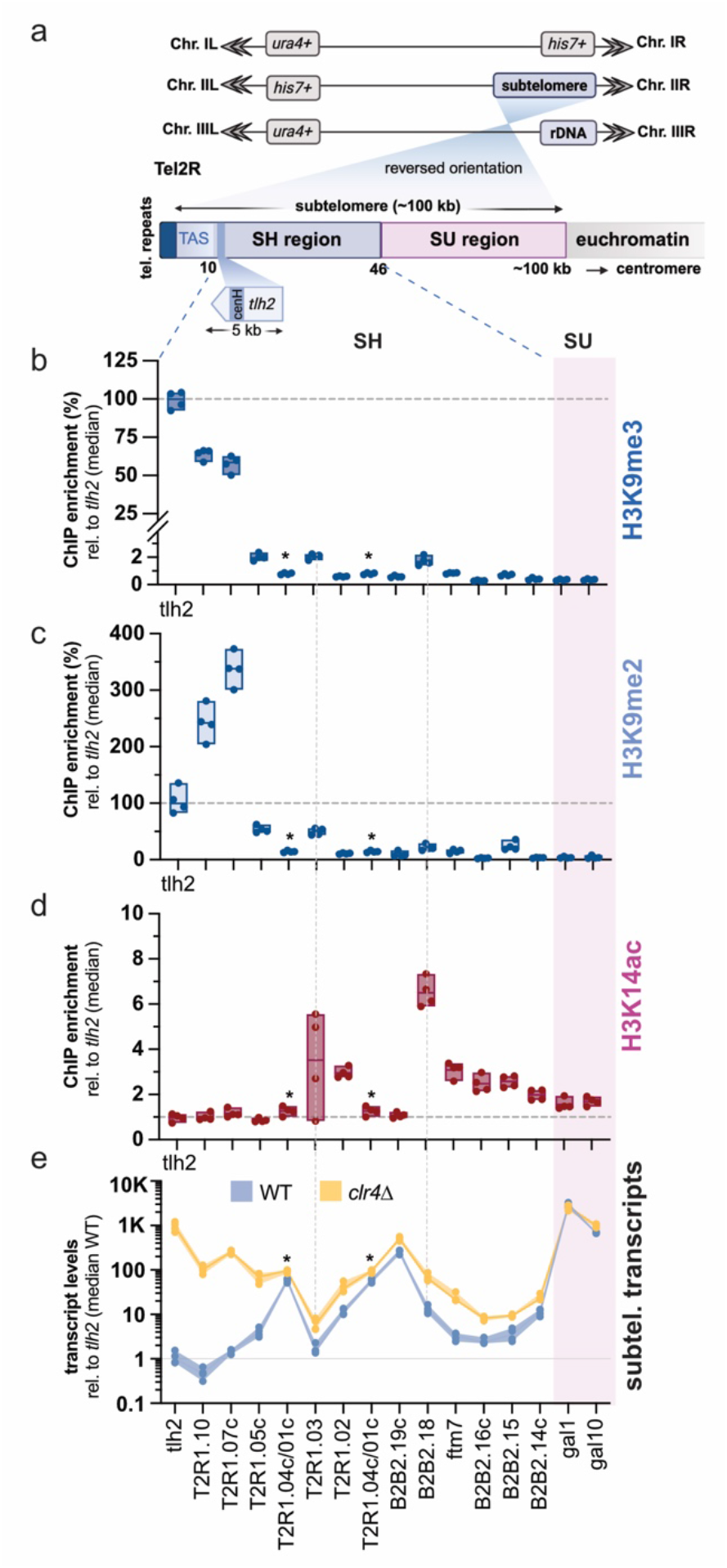
The SH domain carries a heterogeneous chromatin landscape. **(a)** Schematic representation of the *SD4[2R+]* strain with engineered subtelomeric deletions. The 100 kb subtelomeric domain is divided into a ∼50 kb subtelomeric homology (SH, blue) region and a subtelomeric unique (SU, pink) region. All SH sequences were replaced with *ura4* or *his7* markers, leaving only the native subtelomere on the right arm of chromosome II. Following previous studies (Oizumi et al. 2021), the Tel2R arm is displayed in reversed orientation (telomere left, centromere right) to facilitate comparison across subtelomeres and is used consistently throughout this study. Figure created with BioRender. **(b–d)** ChIP–qPCR analysis of histone modifications across genes in the SH region and a portion of the SU region. (b) Enrichment of H3K9me3, (c) H3K9me2, and (d) H3K14ac. Data represent four independent biological replicates (*n* = 4). Input-normalized IP signals were normalized to the mean signal across all subtelomeric loci (pool normalization) and are shown relative to the median tlh2 signal. The SU region is shaded in pink. Primer pairs for amplifying unique target sequences are described in Lei et al 2020. The asterisks denote the gene pair SPBCPT2R1.04c/ SPBCPT2R1.01c displaying 99.9% sequence identity, which cannot be distinguished. **(e)** RT–qPCR analysis of transcript levels from genes in the SH region and a portion of the SU region in wild-type (WT; blue) and *clr4*Δ (yellow) strains. Transcript levels were normalized to *act1* and are shown relative to the median *tlh2* expression level in WT samples to better illlustrate the range of expression levels across the SH region. Independent biological replicates (*n* = 4) are shown as individual data points, with the median and range. The SU region is shaded in pink.

We next assessed whether these chromatin features correlate with transcriptional output by performing quantitative reverse transcriptase (RT-qPCR) assays in *SD4[2R+]* and in a derivative lacking Clr4, the sole H3K9 methyltransferase in *S. pombe*. Within the SH region, downstream of *tlh2*, lie poorly characterized genes that likely arose through gene duplication events and exhibit varying levels of sequence similarity, along with several dubious ORFs (Suppl. Figure S1a). Most SH genes exhibit low transcript levels in the WT background dependent on Clr4, with telomere-proximal genes such as *tlh2* repressed by up to 1000-fold (Figure 1e). The variation in gene expression likely reflects differences in chromatin structure, while also being influenced by intrinsic differences in promoter strength. In contrast, genes in the adjacent SU region display 500-to 1000-fold higher expression that is Clr4-independent (Figure 1e; Suppl. Figure S1b). Notably, two SH genes, *SPBCPT2R1*.*04c* and *SPBB2B2*.*19c*, located ∼28 and ∼38 kb from the telomere, also show high expression and only weak dependence on Clr4, indicating a distinct chromatin context.

To assess whether these features are strain-specific, we compared *SD4[2R+]* to FY1193, a derivative of the commonly used laboratory strain 972 (Ekwall et al. 1999). Although SH sequences vary in DNA sequence and segment copy numbers across different laboratory and wild strains (Oizumi et al. 2021), such variation is primarily confined to the telomere-proximal SH-P region, which is excluded from our analysis. We found that SH gene repression is generally stronger in *SD4[2R+]* than in FY1193, but the overall expression pattern is conserved (Suppl. Figure S1c). These findings indicate that the observed heterogeneity in *SD4[2R+]* is not due to strain-specific variation or the loss of SH sequences on the other chromosome arms.

### Position-specific single-cell reporters identify distinct silencing states across the SH domain

To dissect position-specific heterogeneity in silencing across the SH domain, we developed a single-cell sensor (SCS) reporter system that overcomes the confounding effects of variable promoter strength in endogenous genes and high sequence similarity among the SH genes themselves (Figure 2a). This system builds on our previously established single-cell heterochromatin spreading sensor (HSS) (Greenstein et al. 2018), with specific adaptations for the SH region. It comprises three transcriptionally encoded fluorescent reporters integrated at defined loci in and outside the SH domain, enabling precise, locus-specific quantification of heterochromatinmediated silencing in single cells under a uniform promoter context. To monitor telomere-proximal silencing, we integrated a green fluorescent reporter (superfolder GFP under the *ade6* promoter, *ade6p*, ‘green’) 11 kb from the telomere of *SD4[2R+]*, adjacent to the RNAi-dependent *tlh2* nucleation site (Kanoh et al. 2005). Additional ‘orange’ reporters (monomeric Kusabira Orange 2 under *ade6p*) were inserted at increasing distances (17.5 kb, 28 kb, 33 kb, and 37 kb) to survey nucleation-distal regions. A third reporter (E2Crimson under the *act1* promoter, ‘red’) was integrated at 46.5 kb, marking the SH-SU transition and serving as an internal control to correct for cell-to-cell transcriptional variability caused by intrinsic and extrinsic noise.

**Figure 2:**
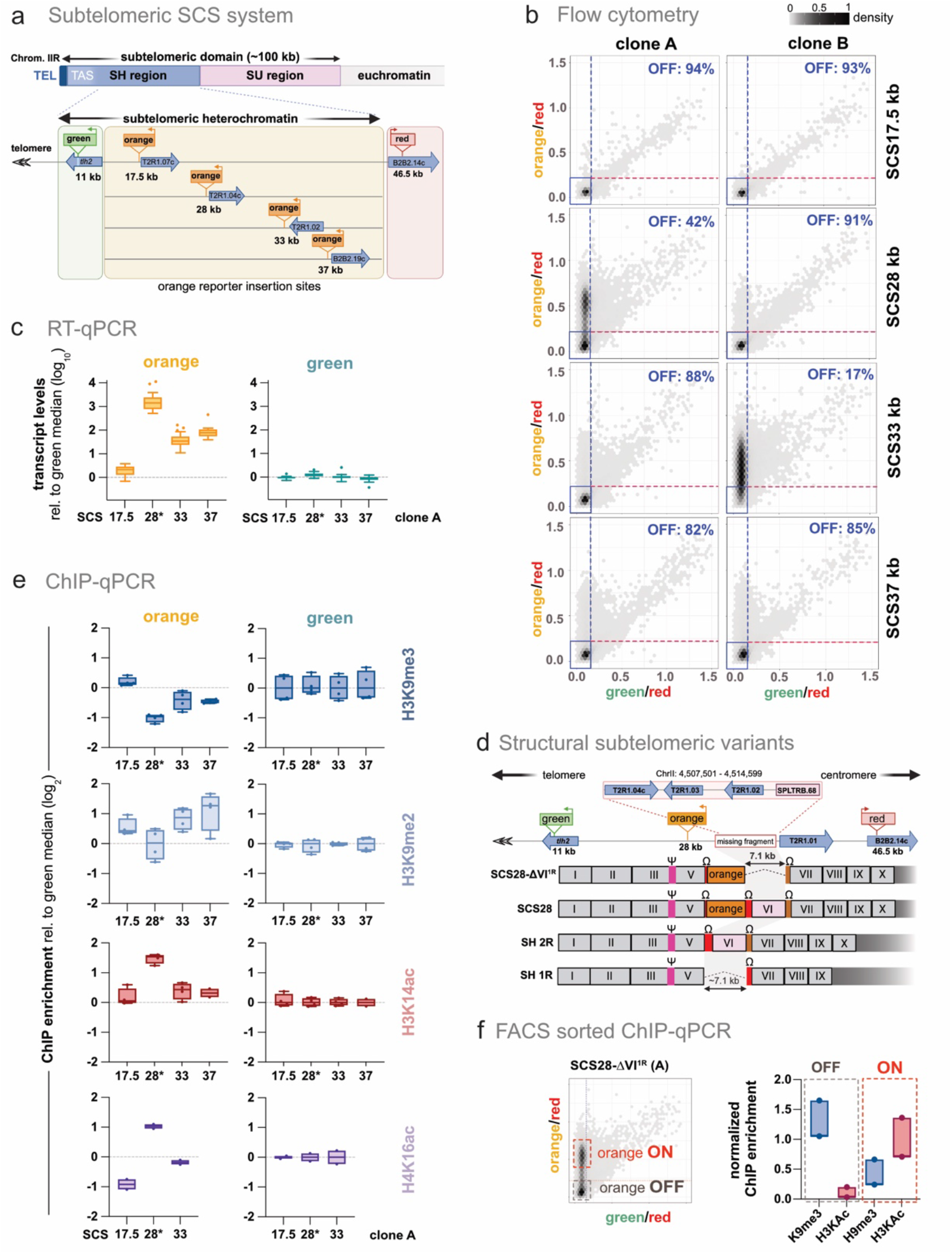
Position-specific single-cell reporters identify distinct silencing states across the SH domain. **(a)** Schematic of the subtelomeric SCS reporter system integrated into the SH region (∼50 kb) of the right arm of chromosome (Chrom. IIR). Each strain contains three reporters: a ‘green’ reporter inserted at 11 kb within the *tlh2* ORF, a ‘red’ reporter at 46.5 kb close to SPBPB2B2.14c, and an ‘orange’ reporter inserted at one of four distinct positions within the SH region (17.5 kb, 28 kb, 33 kb, or 37 kb). The positions of ‘green’ and ‘red’ reporters are consistent across all constructs. Reporter orientation is indicated by arrows, and adjacent genes are labeled. Figure created with BioRender. **(b)** Two-dimensional hexbin density plots showing red-normalized green and orange fluorescence for different SCS reporters. The x-axis represents green fluorescence normalized to red, and the y-axis represents orange fluorescence normalized to red. OFF-state thresholds for ‘green’ and ‘orange’ reporters are indicated by blue and red dashed lines, respectively. Data from two clones, A (left) and B (right), are presented for all SCS constructs. The scale bar at the top right indicates relative cell density, with bin cutoffs plotted as fractions of the most populated bin. **(c)** RT–qPCR analysis of transcript levels from ‘orange’ (left) and ‘green’ (right) reporters in the SCS reporter constructs. Transcript levels were normalized to *act1* and are presented relative to the ‘green’ reporter median expression level. Data from clones A is shown. Data are represented as box-whisker plots (Tukey) from independent biological replicates with *n* = 20 (SCS17.5-A), 16 (28-A*), 29 (33-A), and 18 (37-A). The asterisk in SCS28-A* denotes a 7.1 kb deletion next to the reporter insertion (see text for details). **(d)** Subtelomeric SH regions of Tel1R and Tel2R are depicted based on common block sequences (modified from Oizumi et al. 2021). A 7.1 kb fragment corresponding to SH common block VI is absent in both the naturally occurring SH1R structural variant of chromosomal Tel1R (bottom) and the SCS28 clone A reporter (top), which is hereafter referred to as SCS28-DVI1R. Ψ and Ω homologous box sequences, which share high sequence similarity, are indicated by pink, red, and brown boxes, respectively. The reporter insertion within the Ω sequence is marked by an orange box. Figure created with BioRender. **(e)** ChIP–qPCR analysis of histone modifications at ‘orange’ reporter insertion sites in SCS constructs (clone A). Enrichment of H3K9me3, H3K9me2, H3K14ac, and H4K16ac (from top to bottom). Input-normalized IP signals were normalized to the mean signal across various subtelomeric loci (see Methods for detail) and are shown relative to the median ‘green’ reporter signal (log2 scale). Data points and box plots represent independent biological replicates (for all ChIP experiments n = 4, except for H314ac, 37-A: n = 2; for H3K16ac, all reporters: n = 2). **(f)** ChIP-qPCR on flow cytometry-sorted populations of SCS28-DVI1R. Young, single G2 ‘green’OFF cells were sorted into ‘orange’ON and ‘orange’OFF populations (see Methods). The sorted cells were subjected to ChIP-qPCR analysis of H3K9me3 and H3 pan-acetylation. H3K9me3 ChIP enrichment was normalized to pericentromeric cen-dg repeats, whereas H3 pan-acetylation was normalized to act1. Each data point represents the mean of three technical ChIP replicates from one of two independent biological experiments (n = 2).

We quantified silencing across thousands of cells during logarithmic growth by flow cytometry, analyzing two independent clones for each reporter (A and B; see Methods). Fluorescence signals from ‘green’ and ‘orange’ reporters were normalized to the ‘red’ control and scaled to the fully derepressed state in *clr4Δ* cells, as described previously (Greenstein et al. 2018). Hexbin plots of normalized fluorescence intensities revealed non-uniform silencing distributed across the SH region, consistent with the transcriptional heterogeneity of endogenous genes (Figure 1). The ‘green’ and ‘or-ange’ reporters at 11 kb and 17.5 kb, respectively, exhibited strong silencing in both clones, with over 90% of cells in the OFF states, consistent with their proximity to the *tlh2* nucleation site (Figure 2b). At 28 kb, silencing became heterogeneous, particularly in clone A, which displayed a bimodal distribution of transcriptionally silenced and active cells. We also observed clone-specific differences at 33 kb, whereas silencing at 37 kb was robust in both clones (82–85%), though weaker than at telomere-proximal sites. To validate the chromatin-dependent behavior of the reporters, we measured transcript levels by RT-qPCR (Figure 2c). Consistent with flow cytometry data, the ‘orange’ reporter at 17.5 kb shows low expression similar to the ‘green’ reporter at 11 kb (Figure 2c). The ‘orange’ reporter at 28 kb (clone A) is ∼1000-fold more active, and reporters at 33 kb and 37 kb show intermediate expression levels (∼50–100-fold above green). Conversely, transcript levels of the ‘green’ reporter are not affected by the varying insertion positions of the ‘orange’ reporter. Similarly, while the ‘red’ reporter is located close to the silent SH region, its expression is robust and largely unaffected by the presence of heterochromatin (Suppl. Figure S2a). These findings demonstrate that our position-dependent reporters accurately capture the spatial heterogeneity of subtelomeric gene silencing with greater resolution than endogenous gene readouts.

To test whether reporter insertion perturbs local gene expression, we profiled the transcript levels of endogenous genes. Overall expression patterns were preserved across reporter strains in both wild-type and *clr4*Δ backgrounds (Suppl. Figure S2b), with only minor changes for genes near the reporter insertion sites. However, in SCS28 clone A, three genes adjacent to the 28 kb reporter (*SPBCPT2R1*.*04c, SPBCPT2R1*.*03*, and *SPBCPT2R1*.*02*) were completely absent. These genes belong to a common sequence block (VI) flanked by two nearly identical Ω homologous box sequences (Figure 2d). This 7.1 kb sequence is also naturally absent from the Tel1R arm of the reference laboratory strain 972, likely due to a historical recombination event between the Ω elements (Oizumi et al. 2021). While this structural variant may contribute to the altered silencing profile in clone A, additional evidence indicates that epigenetic mechanisms are also involved (see below). To reflect its structural similarity to the truncated Tel1R chromosome arm, we hereafter refer to clone A as SCS28-ΔVI^1R^.

We next examined histone modifications at the reporter loci using ChIP-qPCR. All reporters display high H3K9me2 and H3K9me3 levels, comparable to those at pericentromeric *cen-dg* repeats (Suppl. Figure S2c). However, we observed variation among the ‘orange’ reporters consistent with the expression data from flow cytometry (Figure 2b) and RT-qPCR (Figure 2c), although differences in these repressive chromatin marks are considerably less pronounced than the corresponding differences in expression levels. In particular, the ‘orange’ reporter in SCS28-ΔVI^1R^, which is expressed more than 100-fold higher than reporters at other sites, exhibits only a twofold reduction in H3K9 methylation. By contrast, H3K14ac levels are markedly increased at the ‘orange’ reporter in SCS28-ΔVI^1R^ (Figure 2c). Consistent with this, deletion of the H3K14 histone acetyltransferase Mst2 reduces ‘orange’ expression in SCS28-ΔVI^1R^ and, to a lesser extent, in SCS33 and SCS37 (Suppl. Figure S3a,b), in agreement with previous reports implicating Mst2 in subtelomere silencing (Gómez et al. 2005). H4K16ac shows a similar pattern, consistent with its enrichment at several active promoters and its reported role in antagonizing heterochromatin spreading (Wang et al. 2013). Because reporter expression is bimodal in SCS28-ΔVI^1R^, these elevated histone acetylation marks may also reflect a bimodal chromatin state; however, it remained unclear whether acetylation increased uniformly across all cells or specifically in cells with high (ON) ‘orange’ expression. To address this question, we used flow cytometry to sort ON and OFF ‘orange’ cell populations, followed by ChIP-qPCR to quantify H3K9me3 and H3 tail acetylation.These experiments revealed that H3K9me3 is reduced in ON cells relative to OFF cells, consistent with a weakened heterochromatic state. In contrast, H3 tail acetylation is nearly absent in OFF cells while it increases dramatically in ON cells (Figure 2f). Together, these results demonstrate that reporter activity is associated with distinct chromatin states, with histone acetylation correlating most closely with the gene expression state.

### Silencing of telomere-distal SH regions is maintained independently of common nucleation pathways

Next, we examined whether previously defined SH heterochromatin nucleation and silencing pathways impact our SCS reporters. We deleted components of the RNAi- and shelterin-dependent pathways, as well as the inner nuclear membrane protein Lem2 (Suppl. Figure S4a). Loss of Tas3, which acts as a scaffold within RITS, the shelterin component Ccq1, or the nuclear membrane protein Lem2 results in modest derepression of the ‘green’ and most ‘orange’ reporters, consistent with their overlapping roles at sub-telomeres (Figure 3a). Despite these similarities, each mutant displays a distinct phenotype: Consistent with the localized and redundant roles of RNAi and shelterin in heterochromatin nucleation, *tas3*Δ exhibits a subtle expression increase of the telomere-proximal 17.5 reporter, whereas *ccq1*Δ displays stronger and broader defects due to additional shelterin binding sites within the SH region (Kanoh et al. 2005; Barrales et al. 2016). In contrast, *lem2*Δ primarily affects telomere-distal silencing, consistent with the broader role of Lem2 in subtelomeric heterochromatin maintenance. By comparison, deletion of the HDAC Clr3, a component of SHREC redundantly recruited by Ccq1 and Lem2 (Barrales et al. 2016; van Emden et al. 2019; Sugiyama et al. 2007), leads to full derepression at every locus (Figure 3a; Suppl. Figure S4b). Intriguingly, whereas the ‘orange’ reporter in SCS28-ΔVI^1R^ displays a bimodal repression pattern in WT cells, this phenotype is suppressed when deleting *tas3* and *ccq1*, resulting in reduced ‘orange’ expression. This suggests that loss of RNAi and shelterin redirects silencing factors from regions dependent on these pathways, such as telomere-proximal and pericentromeric loci, to the ‘orange’ reporter in SCS28-ΔVI^1R^.

**Figure 3:**
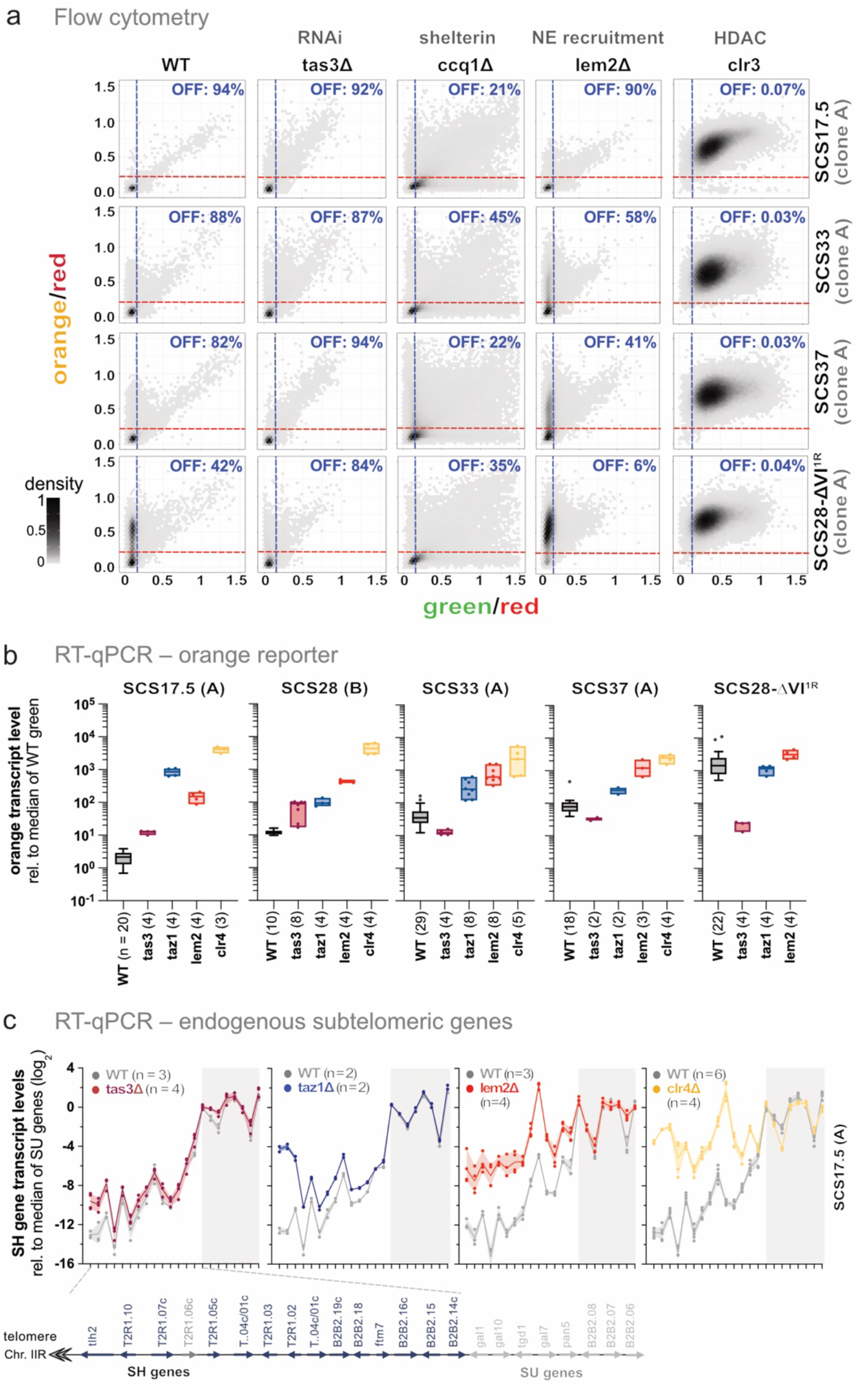
Silencing of telomere-distal SH regions is maintained independently of common nucleation pathways. **(a)** Two-dimensional hexbin density plots showing red-normalized green (x-axis) and orange (y-axis) fluorescence for SCS reporters in various genetic backgrounds. Blue and red dashed lines indicate OFF thresholds for green and orange reporters, respectively. (see Figure 2 for details). Columns represent wild type (WT), *tas3Δ, ccq1Δ, lem2Δ*, and *clr3Δ*. Rows correspond to SCS constructs with variable ‘orange’ reporter positions (17.5, 33, or 37 kb) in the full-length SH-2R region and SCS28-DVI^1R^. Data for WT reporter strains are identical to those shown in Figure 2b. Experiments with *tas3*Δ reporters were conducted independently, with the corresponding WT controls shown in Supplementary Figure S3a. **(b)** RT–qPCR analysis of transcript levels from the ‘orange’ reporters in SCS strains across different genetic backgrounds. Data are shown for reporters located at 17.5, 28, 33, and 37 kb in the SH-2R configuration, and for 28 also in the 1R configuration (SCS28-DVI^1R^). Tested backgrounds include wild type (WT), *tas3Δ, taz1Δ, lem2Δ*, and *clr4*Δ. Individual clones (A, B) are indicated in brackets. Transcript levels were normalized to *act1* and are shown relative to the median ‘green’ reporter expression in WT samples. Data are presented as box-and-whisker plots (Tukey; *n* > 10); ad the number of independent biological replicates is indicated below each plot. **(c)** RT–qPCR analysis of transcript levels for endogenous genes within the SH and SU region across genetic backgrounds in SCS17.5 (clone A). Expression profiles are shown for wild type (WT) and *tas3Δ, taz1Δ, lem2Δ*, and *clr4Δ* and *tas3Δ, taz1Δ*, and *tas3Δ taz1Δ*, with WT values plotted in grey for comparison. Transcript levels were normalized to *act1* and are presented relative to the median of SU-region genes in WT (log_2_ scale). Normalization to the median SU-region expression provides a robust internal reference. Independent biological replicates are shown as individual data points, together with the median and range. The number of independent biological replicates is indicated for each panel. The schematic (below left panel) shows the organization of genes in the SH and SU regions; grey-marked SH genes were excluded from analysis, and all SU-region genes are shown in grey.

We confirmed these position-specific requirements by RT-qPCR (Figure 3b). When examining the different reporters, some common and unique behaviors emerged: deletion of *clr4* abolishes silencing at every reporter, confirming that the observed expression patterns in wild-type cells reflect distinct heterochromatin states rather than sequence-dependent effects. Conversely, deletion of *tas3* predominantly affects telomere-proximal reporter sites. At 17.5kb, transcript levels increase modestly (∼10-fold), consistent with our flow cytometry data and previous findings (Hansen et al. 2006; Kanoh et al. 2005). However, reporters at 33 and 37 kb remain unchanged or show slightly enhanced repression. We observed a similar pattern following deletion of *ago1*, which encodes the siRNA-binding Argonaute subunit of RITS (Suppl. Figure S4c). Notably, the high expression levels observed in WT SCS28-ΔVI^1R^ are reduced by ∼100-fold in *tas3*Δ or *ago1*Δ, reaching values comparable to those in the corresponding non-truncated SCS28 strain (Figure 3b; Suppl. Figure S4c). Loss of *taz1* also leads to strong derepression of the 17.5 kb ‘orange’ reporter and a moderate increase at telomere-distal reporters. Remarkably, although RNAi and Taz1 have been reported to act redundantly at subtelomeric loci (Kanoh et al. 2005; Barrales et al. 2016), their combined loss does not increase but rather further represses ‘orange’ expression in SCS28-ΔVI^1R^ compared to the *taz1*Δ single mutant (Suppl. Figure S3d). Deletion of *lem2* elevates orange expression across reporters, whereas single-cell data obtained by flow cytometry indicates a stronger effect at distal reporters (Figure 3b). This difference may indicate that a subset of cells contributes disproportionately to the RT-qPCR signal. The position-specific functions of Tas3, Taz1, and Lem2 also apply to endogenous subtelomeric SH genes, both in the presence and absence of reporter integration (Figure 3c, top panels; Suppl. Figure S4e). Together, these findings indicate that the contributions of RNAi and shelterin to subtelomeric silencing are largely restricted to telomere-proximal regions.

### Common block sequences locally control gene silencing at the subtelomere

The observation that heterochromatin factors do not regulate gene expression uniformly across the subtelomere, with nucleation factors exerting stronger effects toward the telomere end, raises the question of how local silencing is controlled. Two general models could account for this: (1) silencing is determined primarily by distance from the nucleation site; or (2) silencing is controlled by local *cis-*acting elements. To distinguish between these models, we assessed the impact of deleting two common block sequences, VI and V (Figure 4a, b).

**Figure 4:**
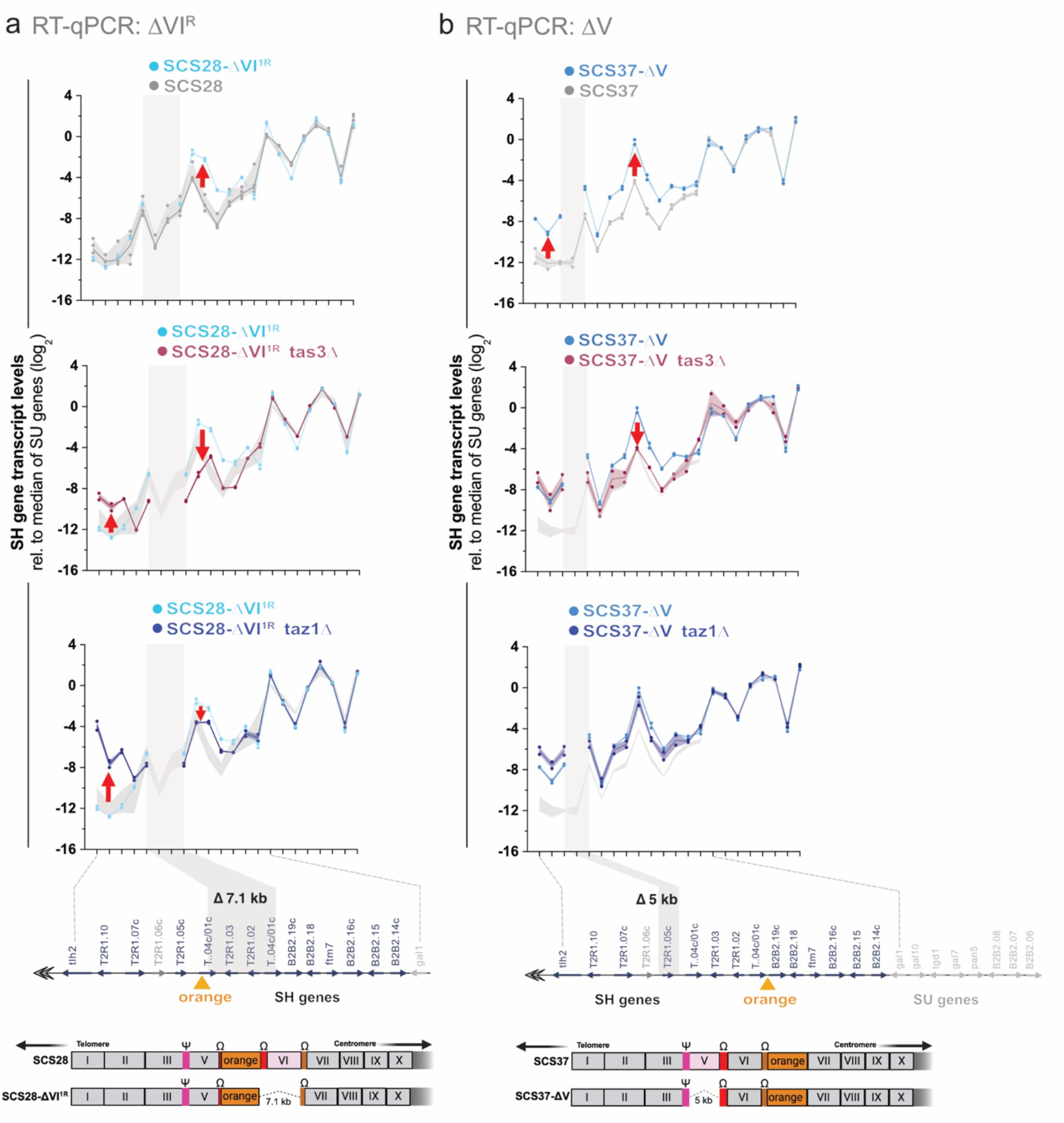
Common block sequences locally control gene silencing at the subtelomere. **(a)** RT-qPCR analysis of the effect of the 7.1-kb block VI deletion on endogenous subtelomeric gene expression. Top: SCS28 SH-2R (SCS28; grey) versus the SH-1R-like block VI deletion strain (SCS28-DVI^1R;^ light blue). Middle: SCS28-DVI^1R^ (light blue) versus SCS28-DVI^1R^ *tas3*D (dark red). Bottom: SCS28-DVI^1R^ (light blue) versus SCS28-DVI^1R^ *taz1*D (dark blue). Two biological replicates, from a representative experiment are shown as individual data points, together with the mean and range, except for SCS28-DVI^1R^, WT: *n* = 4. **(b)** RT–qPCR analysis of the effect of the 5-kb block V deletion on endogenous subtelomeric gene expression. Top: SCS37 SH-2R (SCS37; grey) versus the block V deletion strain (SCS37-ΔV; medium blue). Middle: SCS37-ΔV (medium blue) versus SCS37-ΔV *tas3*Δ (dark red). Bottom: SCS37-ΔV (medium blue) versus SCS37-ΔV *taz1*Δ (dark blue). Two biological replicates are shown as individual data points, together with the mean and range. For (a) and (b), transcript levels were normalized to *act1* and are shown relative to the median expression of SU-region genes (log_2_ scale). The shaded area in the middle and bottom panels represents the data range of the corresponding non-deletion strain. Red arrows indicate loci with substantial expression differences between the corresponding strains. Schematics below the graphs indicate the positions of the reporter insertions and the corresponding block VI or V deletions.

We first focused on the structural variant SCS28-ΔVI^1R^, which lacks the 7.1 kb block VI and shows weaker silencing of the ‘orange’ reporter than its non-truncated sister clone (Figure 2). We examined endogenous gene expression in both reporter strains by RT-qPCR. Consistent with the position-specific effects of RNAi- and shelterin mutants (Figure 3b, c), gene expression near *tlh2* is unaffected by deleting block VI, which lies approximately 15 kb downstream of the RNAi-dependent nucleation site (Figure 4a, top panel). However, several genes downstream of ΔVI (SPBPB2B2.19c, SPBPB2B2.18, and *ftm7*) are moderately derepressed (∼10-fold), indicating that decreasing the distance to nucleation sites does enhance their silencing. Conversely, additional deletion of *tas3* partially restores silencing of these genes in SCS28-ΔVI^1R^ (Figure 4a, middle panel), mirroring the behavior of the ‘orange’ reporter (SCS28-ΔVI^1R^ *tas3*Δ, Figure 3b). As expected, *tlh2-* proximal genes are upregulated in SCS28-ΔVI^1R^ *tas3*Δ in agreement with Tas3’s role in RNAi-dependent nucleation. We observed a similar pattern in combination with *taz1*Δ, although the rescue of silencing downstream of ΔVI appears weaker (Figure 4a, bottom panel).

Next, we deleted block V, which lies upstream of block VI and spans 5 kb. Since this sequence overlaps with the reporter insertion at 28 kb, we performed the deletion in the SCS37 reporter background (SCS37-ΔV). While we observed derepression of genes both upstream and downstream of ΔV, this effect was more pronounced for *tlh2* and adjacent genes (Figure 4b, top panel). Additional deletion of *tas3* partially restores silencing of genes downstream of ΔV, as observed for ΔVI. In contrast, *tlh2*-proximal genes display similarly elevated expression in SCS37-ΔV and SCS37-ΔV *tas3*Δ, indicating that block V and RNAi contribute similarly to silencing of genes upstream of the deletion (Figure 4b, middle panel). Conversely, while additional deletion of *taz1* does not restore silencing in SCS37-ΔV, deletion of block V partially alleviates the silencing defects of taz1Δ at *tlh2*-proximal genes (Figure 4b, bottom panel). We observed similar effects upon deletion of block V in the parental strain lacking reporter insertions, although the changes in gene expression were weaker and largely restricted to genes upstream of the block V deletion. Nevertheless, combining the block V deletion with either *tas3* or *taz1* partially alleviates the silencing defects, as observed in the reporter strains (Suppl. Figure S5). Overall, these findings argue against a simple distance-dependent model, in which repeated blocks act primarily as passive spacers for heterochromatin spreading. Instead, they indicate that the common blocks function as distinct elements that locally shape subtelomeric silencing, as evidenced by the distinct consequences of deleting either block VI or block V, either alone or in combination with *tas3*Δ or *taz1*Δ.

We next investigated whether specific *cis-*regulatory elements within the common blocks directly regulate local silencing. Adjacent to Ω homologous box sequences, block VI is also flanked by two LTRs, SPLTRB.69 and SPLTRB.68 (Suppl. Figure S6a); the latter is absent in SCS28-ΔVI^1R^. Because LTRs have been proposed to act as boundary elements separating euchromatin from heterochromatin domains (Steglich et al. 2015; Cam et al. 2005), we hypothesized that the loss of SPLTRB.68 might underlie the altered silencing phenotype in SCS28-ΔVI^1R^ . To test this, we deleted SPLTRB.68 using CRISPR in the non-truncated sister clone SCS28 and another reporter strain, SCS33. Neither deletion substantially affects ‘orange’ expression (Suppl. Figure S6b), indicating that loss of SPLTRB.68 alone is insufficient to explain the phenotype. However, removing a different LTR (SPLTRB.67) located further downstream caused a >10-fold increase in ‘orange’ expression relative to the corresponding WT strain (Suppl. Figure. S6b). These findings suggest that while SPLTRB.68 is largely dispensable for repression, specific LTR elements, such as SPLTRB.67, can modulate local silencing and contribute to the organization of SH heterochromatin.

Having found only limited support for a role of LTRs in this phenotype, we next asked whether common block sequences also harbor heterochromatin nucleation sites. SH genes adjacent to the 7.1 kb fragment in SCS28-ΔVI^1R^ maintain silencing independently of RNAi and the shelterin complex (Figure 4a), suggesting that this segment or nearby sequences may function as a cryptic heterochromatin nucleation site capable of recruiting Clr4, independently of these nucleation pathways and pre-existing H3K9 methylation. To identify such sites, we examined Clr4 enrichment in an *ago1*Δ *H3K9R* strain at subtelomeric regions and detected a small but distinct Clr4 peak near the 7.1-kb segment within block V (Suppl. Figure S6c and S6d). Comparable peaks are present on both Tel1L and Tel2R at similar, though not identical, positions. Intriguingly, the Clr4 binding site on Tel2R is adjacent to H3K9me3 peaks that persist in Swi6^*HP1*^ mutants deficient in heterochromatin spreading (Suppl. Figure S6e) (Kennedy et al. 2024). These Clr4 and H3K9me3 peaks overlap with or are close to the ncRNAs SPNCRNA.2012, SPNCRNA.6640, and SPNCRNA.6641. Similar ncRNAs have been linked to RNAi-independent Clr4 recruitment at the pericentromere and heterochromatin islands (Khanduja et al. 2024; Sugiyama et al. 2016; Zofall et al. 2012; Vo et al. 2019; Hiriart et al. 2012). These findings suggest that this telomere-distal region may harbor cryptic, ncRNA-dependent nucleation sites.

Together, these data indicate that common block sequences define autonomous subtelomeric silencing domains rather than acting as passive spacers for heterochromatin spreading. Telomere-proximal regions remain dependent on RNAi- and shelterin-mediated nucleation, whereas more distal regions may harbor cryptic, potentially ncRNA-dependent nucleation sites and are further modulated by LTR elements.

### Silencing at distal subtelomeric subdomains has unique genetic requirements

The phenotypic differences among core pathway mutants support the idea that silencing at each locus relies on distinct genetic requirements, potentially involving distinct sets of *trans-*acting factors operating across the subtelomeric region. To explore this further, we performed a targeted screen using the SCS28-ΔVI^1R^ and SCS37 reporters, which represent different structural variants and display distinct epigenetic behaviors. SCS28-ΔVI^1R^ additionally has the advantage that it allows the detection of silencing antagonists. We crossed these reporters to a curated set of chromatin factor mutants (partial list from Greenstein et al., 2022; see Supplemental Tables S4-S6) and examined the resulting strains by flow cytometry (Figure 5a). We prioritized genes that affect the silencing at ‘orange’, as factors leading to de-repression of ‘green’ overlapped with known, general regulators of heterochromatin (Suppl. Figure S7a, b). Because strains in the mutant library carry intact SH regions on all subtelomeric arms, the genetic crosses generated F1 progeny with mixed subtelomeric derivates. Nonetheless, flow cytometry analysis on F1 progeny revealed no detectable differences between these progeny and the parental reporter strains (Suppl. Figure S7c). To quantify silencing phenotypes of the SCS28-ΔVI^1R^ and SCS37 reporters, we analyzed the distribution of individual cells by dividing the population into four zones based on the level of ‘orange’ repression and quantified the fraction of cells in each zone in WT and mutant strains (Figure 5b). For SCS37, in which the ‘orange’ reporter is strongly repressed in WT cells, most mutants exhibit loss of silencing, whereas mutants in SCS28-ΔVI^1R^ display both loss and gain of silencing (Suppl. Figure S7a, b). Using the combined results from both reporter screens, we performed k-means clustering, which identified five major clusters (Suppl. Figure S7d). Because two clusters displayed highly similar phenotypes and also showed enrichment for members of the same protein complexes, we combined them into group II, comprising subgroups IIa and IIb. Group I contains mutants with enhanced silencing primarily in SCS28-ΔVI^1R^. Groups IIa and IIb comprise mutants with silencing defects exclusively in SCS28-ΔVI^1R^. Group III contains mutants with strong silencing loss in SCS28-ΔVI^1R^ and partial silencing defects in SCS37, whereas group IV comprises mutants with pronounced silencing defects of both reporters.

**Figure 5:**
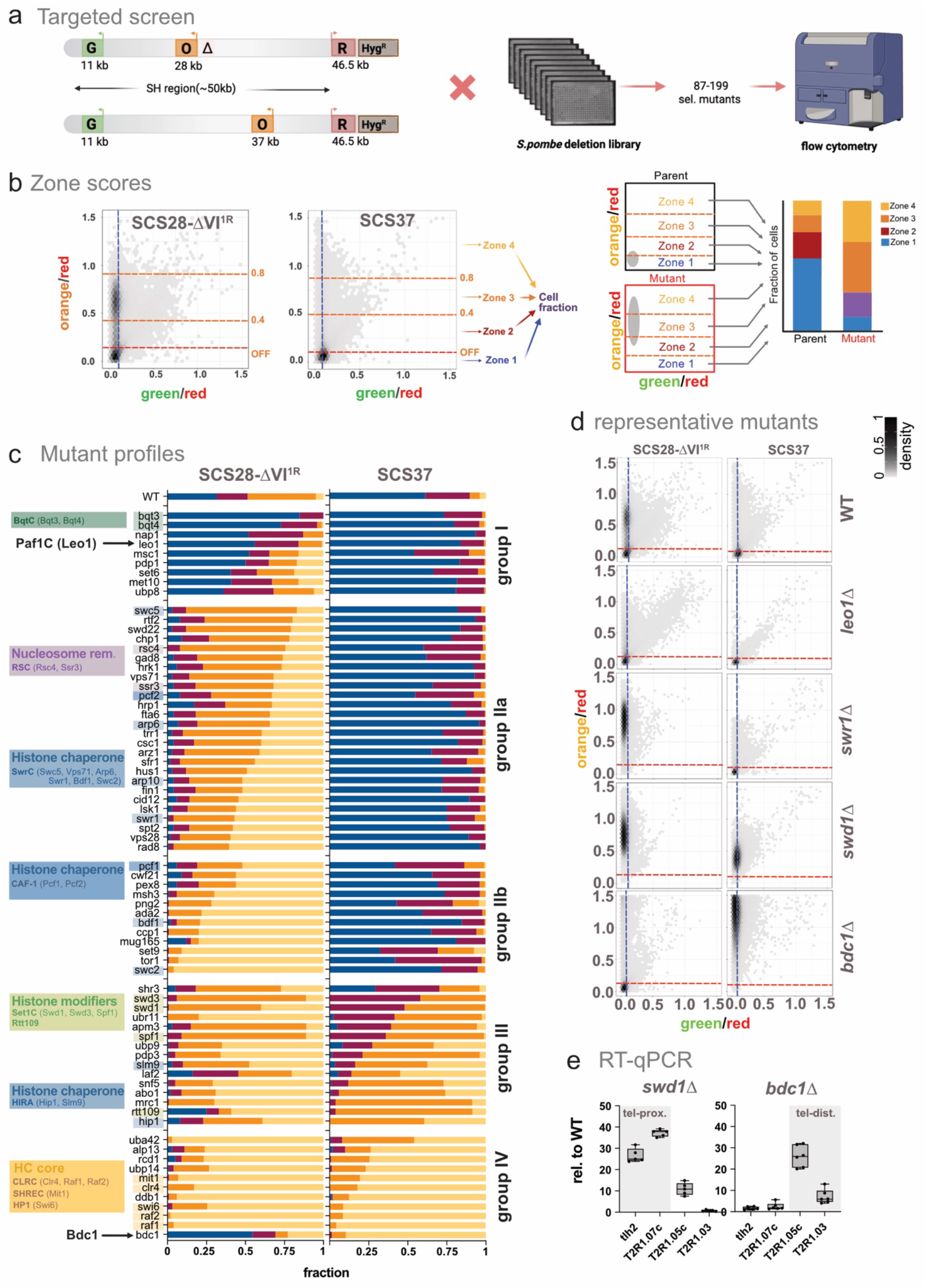
Silencing at distal subtelomeric subdomains has unique genetic requirements. **(a)** Schematic illustrating the fluorescence-based targeted screen. SCS28-DVI^1R^ and SCS37 reporter constructs contain orange reporters inserted at 28 kb or 37 kb, respectively, with ‘green’ and ‘red’ reporters fixed at 11 kb and 46.5 kb. Both reporters were crossed with a deletion library of ∼3000 non-essential genes. A subset of 87–199 mutants were selected for FC analysis. **(b)** Two-dimensional hexbin density plots showing green (x-axis) and orange (y-axis) fluorescence normalized to red for SCS reporter constructs. Blue and red dashed lines indicate OFF thresholds for ‘green’ and ‘orange’ reporters, respectively, which are based on fluorescence in either channel in a ‘red’ only strain (see Methods). Orange fluorescence was further stratified into four zones (zone 1–4) based on increasing orange/red ratios using cutoffs at the OFF threshold, 0.4, and 0.8. The fraction of cells within each zone was quantified to assess the distribution of ‘orange’ reporter expression. This zonal classification was applied to identify mutants that promote activation of the orange reporter, as indicated by increased cell fractions in Zones 3 and 4. **(c)** Stacked bar plots showing the distribution of cells across four fluorescence expression zones (zone 1–4, increasing orange fluorescence normalized to red) in wild-type and mutant strains in the SCS28-DVI^1R^ (left panel) or SCS37 (right panel) reporter. The shared set of genes identified in both screens was grouped into five clusters based on zone distribution using optimal k-means clustering (k = 5; see Supplementary Figure S7d). Because two clusters displayed highly similar phenotypes and shared members of the same protein complexes, they were combined into group II (subgroups IIa and IIb) for subsequent analyses. Each horizontal bar represents one mutant strain. For each group, mutants are ordered top to bottom by the increasing fraction of cells in zones 3 and 4 in the SCS28-DVI^1R^ screen (left). Corresponding zone distributions for the same mutants are shown in the SCS37 screen (right). Wild-type distributions are included at the top for comparison. Gene names are annotated on the left and grouped by functional categories, including transcriptional regulators (e.g., Paf1C), chromatin remodelers (e.g., SwrC, RISC), histone chaperones (e.g., CAF-1, HIRA), histone modifiers (e.g., Set1C, Rtt109), and heterochromatin core components (e.g., CLRC, SHREC, HP1). This classification facilitates systematic identification of regulators influencing silencing or derepression of the orange reporter. **(d)** Two-dimensional hexbin density plots showing green (x-axis) and orange (y-axis) fluorescence normalized to ‘red’ for SCS28-DVI^1R^ and SCS37 reporter constructs. Wild-type profiles are shown at the top for reference. Representative mutants are displayed below. The left panel shows wild-type and mutants in the SCS28-DVI^1R^ background; the right panel shows wild-type and mutants in the SCS37 background. **(e)** RT-qPCR analysis of endogenous gene expression in independently generated *swd1* and *bdc1* knockout strains in the parental *SD4[2R+]* background lacking the reporters. Transcript levels were normalized to *act1* and are shown relative to the median WT value. Five biological replicates from three independent experiments are shown as individual data points in box-and-whisker plots, except *bdc1* knock out strains: *n* = 6. Shaded areas highlight telomere-proximal (*tlh2*, SPBCPT2R1.07c; left panel) and telomere-distal genes (SPBCPT2R1.05c, SPBCPT2R1.03, right panel).

Noteworthy examples from group I (mutants that enhance silencing of SCS28-ΔVI^1R^) include the Paf1C subunit Leo1, which also partially enhances silencing in SCS37 (Figure 5c). This finding is consistent with previous reports that revealed a role of Paf1C in promoting nucleosome turnover while antagonizing heterochromatin assembly and spreading (Verrier et al. 2015; Kowalik et al. 2015; Sadeghi et al. 2015). Additional factors include the NE-bound proteins Bqt3 and Bqt4 implicated in telomere tethering (Figure 5c). We also identified members of Ino80C (Ies2, Ies4, Iec1 Iec3) and two jumonji proteins, Jmj2 and Jmj4, which are potentially involved in H3K4me3 methylation (Huarte et al. 2007) (Suppl. Figure S7a). This may be related to the formation of H3K4me-dependent boundaries in open chromatin (see discussion).

Group II (subgroup IIa and IIb) contains mutants with silencing defects in SCS28-ΔVI^1R^. Among those mutants, chromatin remodeling factors display unusual behaviors. Previously, mutations in these genes have been implicated in restricting heterochromatin spreading (Greenstein et al. 2022) and assembly (Sahu et al. 2024). Surprisingly, we found that several remodeling complex members are required for ‘orange’ silencing in SCS28-ΔVI^1R^ but not SCS37, including Swr1C, which is involved in H2A.Z deposition (Swr1, Swc2, Swc5, Arp6, Vps71, Bdf1), and RSC (Rsc4 and Ssr3). We also found that members of the histone chaperones CAF1 (Pcf1, Pcf2) specifically affect SCS28-ΔVI^1R^. This behavior contrasts with that at the MAT locus, likely reflecting the role of SH as a buffering sink for free heterochromatin factors (Murphy and Berger 2023; Tadeo et al. 2013; Tashiro et al. 2017). In this scenario, the gain of silencing observed in remodeler mutants at pericentromeric and MAT loci comes at the expense of factors lost from the SH region, with the SCS28-ΔVI^1R^ reporter exhibiting a much broader sensitivity to remodeling complex members.

Group III mutants exhibit silencing loss at ‘orange’ in both reporters but have a stronger effect in SCS28-ΔVI^1R^. Besides the replication fork-associated protein Mrc1 previously shown to promote heterochromatin inheritance (Charlton et al. 2024; Toda et al. 2024; Yu et al. 2024; Kawakami et al. 2024), this group is particularly enriched for members of the Set1C complex (Figure 5c, d) and subunits of the HIRA histone chaperone (Hip1, Slm9; Figure 5c; Suppl. Figure S7e). We validated this telomere-proximal biased silencing effect for the Set1C member Swd1 and the HIRA subunit Hip1 by RT-qPCR (Figure 5e; Suppl. Figure S7f). While several of these group III genes have been reported to affect heterochromatin silencing (Yamane et al. 2011; Muhammad et al. 2024; Lorenz et al. 2014), mutants in these particular complexes overlap with those we previously demonstrated to specifically disrupt ectopic heterochromatin at the euchromatic *ura4* locus, while having a weaker impact on constitutive sites (Greenstein et al. 2022).

Group IV mutants that disrupted ‘orange’ silencing in both reporters include several previously characterized heterochromatin factors, including the Clr6 deacetylase complex (Greenstein et al., 2022), the E3 ubiquitin ligase adapter protein Ddb1 (Braun et al. 2011), and members of the H3K9 methyltransferase Clr4 complex and the HP1 homolog Swi6.

Beyond the factors conforming to groups I-IV, we identified several mutants with distinct phenotypes. The most striking example is *bdc1*, which encodes a bromodomain protein of the NuA4 HAT complex. Bdc1 antagonizes silencing at 28 kb but is essential for silencing at 37 kb. We also confirmed its requirement for repression of telomere-distal genes in the parental *SD4[2R+]* strain lacking the reporters (Figure 5e). Bdc1 has not previously been implicated in heterochromatin regulation and may perform a unique, context-dependent role at SH domains. Although too few mutants were specific to SCS37 to define a separate group, one notable example is Rtt109, the H3K56 acetyltransferase in *S. pombe* (Xhemalce et al. 2007). Although these factors do not belong to a single molecular pathway, their distinct phenotypes underscore the modular nature of SH heterochromatin and the diverse regulatory mechanisms acting on individual subdomains.

### SH chromatin undergoes unique dynamic changes not seen at other constitutive heterochromatin loci

Previously, we documented widespread fluctuation in silencing over time at the MAT locus using a sensor positioned 4 kb distal to the constitutive *cenH* nucleation site, even as the sensor at the nucleation site remained silent (Greenstein et al. 2018). To determine whether SH heterochromatin exhibits similar temporal dynamics, we monitored the 28 kb ‘orange’ reporter using the Fission Yeast Lifespan Microdissector (Fig) (Spivey et al. 2017), which enables single-cell tracking of reporter expression over more than 40 generations (Figure 6a). We selected the SCS28-ΔVI^1R^ reporter and its non-truncated SCS28 sister strain due to their distinct silencing ground states. A *clr4*Δ strain served as a control to scale ‘green’ and ‘orange’ fluorescence in the absence of heterochromatin. Live-cell imaging showed that the ‘green’ reporter, located near the *tlh2* nucleation site, remains fully repressed throughout the experiment in both clones, as expected (Figure 6b, top traces). In contrast, as predicted from flow cytometry, the ‘orange’ reporter displays clone-specific behavior: the majority of cells occupy fully or intermediate ON states in SCS28-ΔVI^1R^, whereas those in SCS28 are predominantly OFF. Nonetheless, both clones show moderate switching activity, with infrequent transitions between the ON and OFF states observed in a small number of cells (see example traces in Figure 6b, depicted in grey). Quantification confirmed that the majority of SCS28-ΔVI^1R^ cells remain ON and most SCS28 cells remain OFF for the duration of the experiment, indicating low frequency transitions (Figure 6c).

**Figure 6:**
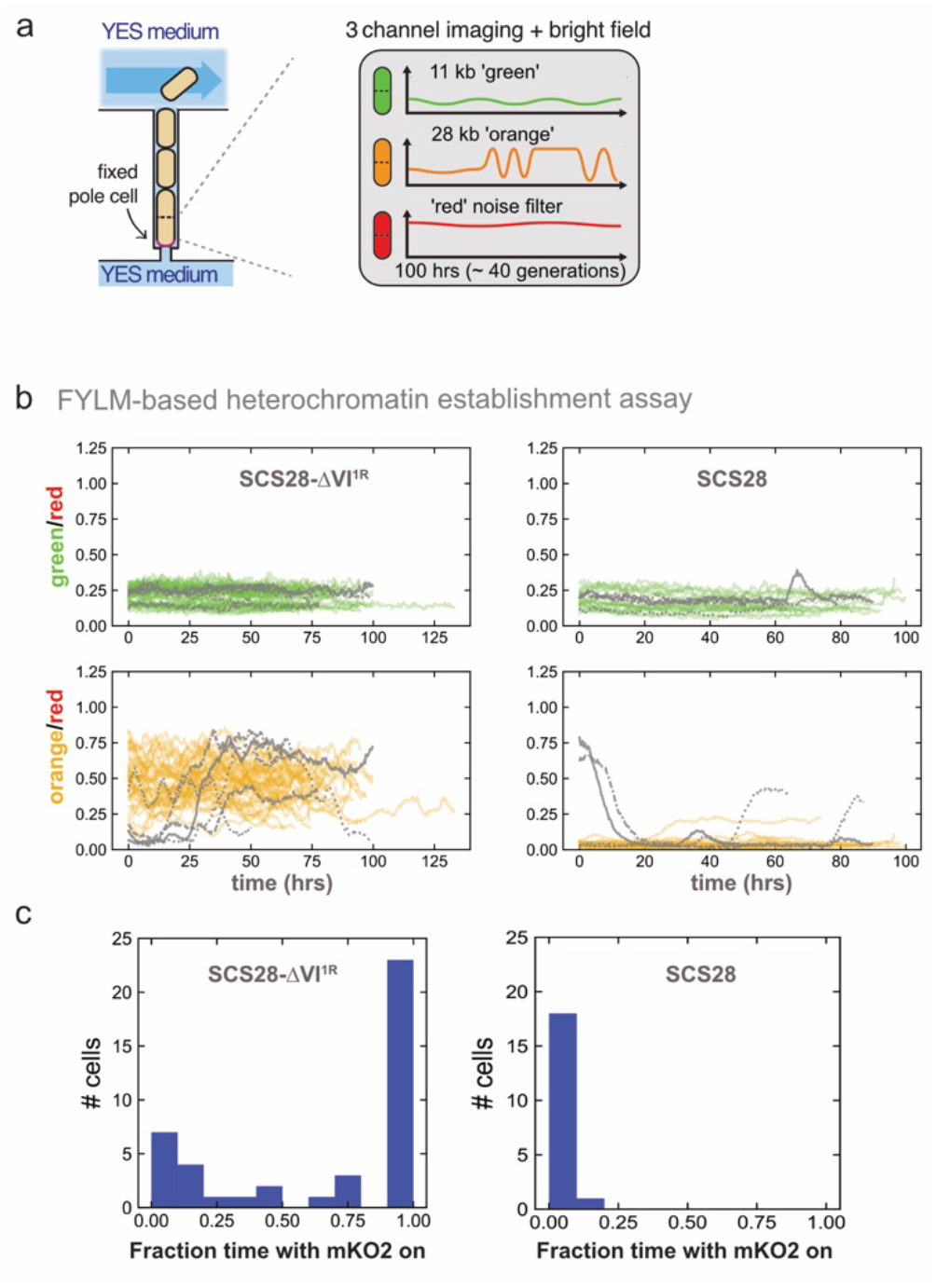
Intergenerational tracking reveals unique dynamic changes in SH chromatin. **(a)** Schematic of FYLM experiment. **(b)** Normalized fluorescence traces for cells that maintained the nucleation sensor (‘green’) in the OFF state throughout an experiment. Thresholds for ON and OFF states were determined within each experiment using signals from *clr4Δ* cells (see Methods). Cells were pooled from three independent experiments (total number: SCS28-DVI^1R^, *n*=42 cells; SCS28, *n*=19 cells). Grey stylized lines indicate cells whose orange-to-red ratio crossed between ON and OFF states during the experiment. **(c)** Histograms of time spent in ‘orange’ ON and OFF states for each cell. For each cell, the number of hours where the cell’s orange-to-red ratio was above the ON threshold was divided by the total number of hours collected for that cell.

The temporal behavior of both 28 kb reporters differs markedly from that observed at the MAT locus (Greenstein et al. 2018). There, we observe either extreme fluctuations at ‘orange’ in all cells, or no fluctuations at all, across the entire time course depending on the heterochromatin nucleator. In contrast, cells carrying the SCS28 reporters remain stably ON or OFF throughout the experiment, with only infrequent fluctuation. These findings demonstrate that SH heterochromatin at the ∼28 kb region, independent of its precise genetic context, adopts a distinct, metastable epigenetic state.

### The SH-domain contains subdomains with unique epigenetic behaviors

Having established metastability at SCS28, we next asked whether the differences in the steady-state distributions among the ‘orange’ reporters (Figure 2) reflect distinct epigenetic states along the subtelomeres. To test this, we used a classic recovery approach, which monitors heterochromatin re-establishment following transient perturbation with the HDAC inhibitor trichostatin A (TSA) (Ekwall et al. 1997). To globally reset heterochromatin states, we cultured our reporter strains (SCS17.5, 28-ΔVI^1R^, 33, and 37) overnight in TSA (Hall et al. 2002; Greenstein et al. 2018), then removed the inhibitor and propagated cells continuously for up to 28 days (Figure 7a). For the 28 kb and 33 kb sensors, we examined both clones (A and B). After TSA withdrawal, we propagated multiple isolates of each sensor strain and monitored silencing dynamics over time. All sensors exhibited complete derepression of the ‘orange’ reporter, while the ‘green’ reporter at 11 kb shows partial derepression among all strains (Figure 7b, Suppl. Figure 8a). For examining re-establishment, we chose day 12 and 28, as silencing profiles had stabilized by that time (Figure 7b, Suppl. Figure 8a-c).

**Figure 7:**
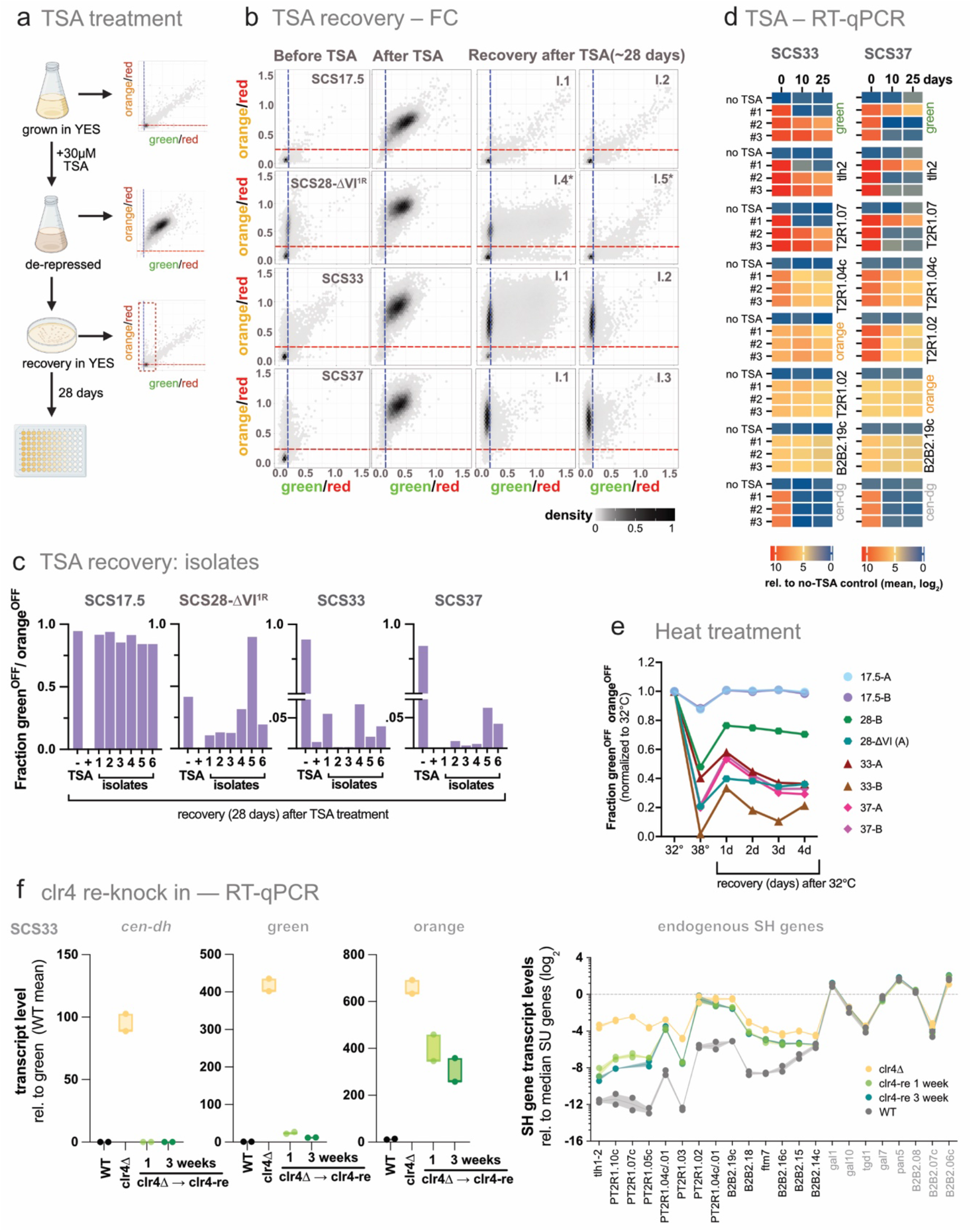
The SH-domain contains subdomains with unique epigenetic behaviors. **(a)** Schematic illustrating trichostatin A (TSA) treatment and recovery assay. **(b)** Two-dimensional hexbin density plots showing green (x-axis) and orange (y-axis) fluorescence normalized to ‘red’ for SCS reporter constructs. Blue and red dashed lines denote OFF thresholds for green and orange reporters, respectively, as in Figure 2b. Columns show fluorescence profiles in untreated wild-type (WT), following treatment with 30 μM TSA, and in two independent isolates after 28 days of recovery in nutrient-rich media. Rows correspond to SCS strains containing ‘orange’ reporters inserted at 17.5, 28, 33, or 37 kb; ‘green’ and ‘red’ reporters are as above. **(c)** Bar plots showing recovery dynamics of SCS reporter clones after TSA treatment. The y-axis indicates the fraction of cells in the ‘orange’ OFF / ‘green’ OFF fluorescence state. Conditions include untreated cells, 30 µM TSA-treated cells, and isolates recovered for 28 days in nutrient-rich medium. Six independent post-recovery isolates are shown for each SCS construct. **(d)** RT-qPCR analysis of the recovery dynamics of reporter and endogenous subtelomeric gene expression in SCS33 and SCS37. Treatment was performed as described in (b), followed by recovery for 0 (no recovery), 10, or 25 days. For each strain, three independent isolates following recovery (#1-3) and a non-treated control (no TSA) were analyzed. The day 0 sample is shared among all three recovery isolates. Transcript levels were normalized to *act1* and are shown relative to the mean expression of the corresponding no-TSA control value for each transcript across days 0, 10, and 25 (log_2_ scale). **(e)** Time course of chromatin state re-establishment following transient disruption. Line plot showing the recovery dynamics of SCS reporter clones after 38°C heat treatment and return to 32°C. The y-axis indicates the fraction of cells exhibiting the ‘orange’ OFF / ‘green’ OFF’ fluorescence state. Timepoints correspond to baseline, immediately after heat treatment, and up to four days of recovery. Each line represents an individual SCS reporter clone (A, B) with an ‘orange’ reporter inserted at 17.5, 28, 33, or 37 kb. **(f)** Transcript-level recovery following *clr4* reintroduction in SCS33 (clone A). **Left panel:** RT–qPCR analysis of transcripts from pericentromeric *cen*-*dg* repeats, the ‘green’ reporter (11 kb), and the ‘orange’ reporter (33 kb). After *clr4* deletion, the gene was reintroduced, and cells were sampled after 1 and 3 weeks of recovery. Transcript levels were normalized to *act1* and shown relative to the mean green reporter expression in WT cells. Each data point represents an independent biological replicate (*n* = 2). **Right panel:** RT–qPCR analysis of transcript levels across the SH region and the SU region in SCS33 (clone A). The line plot shows gene expression in wild-type (grey), *clr4Δ* (yellow), and after *clr4* reintroduction with 1 week (light green) or 3 weeks (dark green) of recovery. Transcript levels were normalized to *act1* and shown relative to the median expression of SU-region genes in wild type (log_2_ scale). Data are from two independent biological replicates, as in the left panel, and shown as mean with range.

In all strains, we observed rapid re-establishment of silencing of the ‘green’ reporter, indicating that *tlh2*-proximal heterochromatin is quickly restored (Figure 7b and quantified in Figure 7c). In contrast, the silencing dynamics of the ‘orange’ reporter vary depending on its genomic position. At 17.5 kb, silencing is uniformly restored, indicating that rapid restoration of *tlh2*-proximal hetero-chromatin extends to this site. However, at 28 kb, recovery is heterogeneous. Isolates derived from the bimodal parental clone SCS28-ΔVI^1R^ adopted a range of states following recovery, from derepressed to fully silenced states. Similarly, TSA-recovered isolates from the non-truncated SCS28 sister clone exhibit both repressed and bimodal silencing profiles, the latter being reminiscent of SCS28-ΔVI^1R^ (Suppl. Figure S8a and quantified in Figure 7c and Suppl. Figure S8b). These results suggest that the region around 28 kb can adopt at least two distinct epigenetic states—fully repressed or bimodal—that can emerge spontaneously and persist over time. This further implies that the variability in SCS28-ΔVI^1R^ is not solely attributable to genetic difference but is instead also influenced by the specific chromatin context at this locus.

The ‘orange’ reporters at 33 and 37 kb behaved distinctly from both 17.5 and 28 kb. In the unperturbed state, both reporters are stably repressed (Figure 2b) except for SCS33-B. However, neither locus fully regained silencing following TSA treatment (Figure 7b, quantified in Figure 7c, Suppl. Figure S8a-c), indicating that regions distal to 28 kb harbor a more fragile epigenetic state. This is also evident at the level of the neighboring endogenous genes, which we tracked by RT-qPCR in SCS33 and SCS37 (Figure 7d). Here, we observed rapid recovery of *cen-dg* in all isolates, whereas ‘green’, *tlh2*, and SPBCPT2R1.07c, which lies downstream of ‘orange’ in SCS17.5, recovered quickly in half the isolates, consistent with flow cytometry (Figure 7b). In contrast, the telomere distal genes SPBCPT2R1.04c, SPBCPT2R1.02, and SPBPB2B2.19c genes as well as ‘orange’ at either 33 kb or 37 kb experienced only weak recovery.

One possible explanation for the lack of recovery at telomere distal sites is that chromatin in this region is more susceptible to the loss of H3K9-methylated nucleosomes. If so, re-establishing its repressed state may require genetic conditions that stabilize H3K9 methylation. Epe1, a known antagonist of heterochromatin, promotes nucleosome turnover at H3K9-methylated regions (Aygun et al. 2013). To test this hypothesis, we repeated the TSA recovery experiment in an *epe1Δ* background. Indeed, silencing of the ‘orange’ reporters at 33 kb and 37 kb was partially restored under this condition (Suppl. Figure 8d), indicating that chromatin at these positions is particularly sensitive to disruption of H3K9 methylation and benefits from protection against Epe1-mediated turnover.

To determine whether this heightened sensitivity extends beyond HDAC inhibition, we next disrupted gene silencing by transient heat stress, an orthogonal perturbation of heterochromatin. We previously found that transient heat exposure weakened heterochromatin (Greenstein et al. 2018), likely through relocalization of Dcr1 from the nucleus to the cytoplasm under elevated temperature (Woolcock et al. 2012). SCS strains were grown at 32ºC, shifted to 38ºC overnight, and then returned to 32ºC to monitor the re-establishment of silencing. Both SCS17.5 clones showed only minimal induction of ‘orange’ expression during temperature treatment and rapidly recovered silencing upon return to 32ºC (Figure 7e). In contrast, the SCS28, SCS33, and SCS37 reporters displayed substantial loss of silencing at 38ºC. Among these, only SCS28-ΔVI^1R^ partially restored silencing, whereas the SCS33 and SCS37 strains failed to re-establish detectable repression. Notably, the 28 kb sensor also re-establishes silencing after TSA treatment, indicating that its capacity for heterochromatin recovery is maintained across distinct perturbations. By contrast, the 33 kb and 37 kb sensors, although robustly silenced under steady-state conditions, are highly sensitive to both heat stress and HDAC inhibition.

Finally, to determine whether the impaired recovery of telomere-distal heterochromatin also extends to de novo heterochromatin establishment, we transiently deleted the *clr4* gene and measured the silencing status of both reporters and endogenous subtelomeric genes in SCS33 (clone A) by RT-qPCR (Figure 7f). Following *clr4* deletion and subsequent re-integration into its endogenous locus using CRISPR, we found that pericentromeric and subtelomeric loci dependent on RNAi quickly re-established silencing, particularly at the *dh* repeats. In contrast, silencing at the ‘orange’ reporter at 33 kb remained incomplete, even 3 weeks after *clr4* reintegration. Analysis of individual subtelomeric genes revealed partial recovery at telomere-proximal loci, whereas more telomere-distal genes located downstream of the reporter insertion site exhibited little or no repression. Together, these findings indicate that de novo heterochromatin establishment is markedly inefficient within the SH domain at telomere-distal loci that are distant from RNAi-dependent nucleation sites.

## Discussion

By deploying a system that tracks a single SH domain in *S. pombe* together with single-cell sensors, we probed both the establishment and epigenetic behavior of SH heterochromatin. Three central findings emerge from this work: (1) Heterochromatin assembles within discrete SH subdomains, possibly through additional cryptic nucleation sites, each exhibiting distinct genetic dependencies and dynamic behaviors; (2) SH heterochromatin displays a spectrum of maintenance profiles across individual cells; (3) SH heterochromatin harbors characteristics that set it apart from constitutive heterochromatin at other genomic loci.

### Formation of SH heterochromatin likely occurs in unique subdomains

Our findings indicate that the genetic requirements for subtelomeric silencing vary substantially across SCS sites, underscoring the position-specific organization of heterochromatin. The RNAi pathway exerts a short-range effect at endogenous SH genes, selectively increasing the expression of the telomere-proximal ‘orange’ reporter, SCS17.5. Shelterin acts over a broader region but has weaker effects on the 28–37 kb reporters and no detectable role near the SH/SU border. Finally, silencing of SCS28-ΔVI^1R^ and SCS37 depends on distinct sets of factors. Notably, silencing of ‘orange’ in SCS28-ΔVI^1R^, which mimics the naturally truncated Tel1R arm lacking the 7.1 kb fragment, is restored in RNAi-deficient mutants (*tas3*Δ, *ago1*Δ) to levels comparable to the SCS28 sister clone, which retains the complete SH sequence.

We propose two models that can explain these observations: Heterochromatin nucleation occurs exclusively via telomere-proximal RNAi and shelterin pathways, with local chromatin structures modulating silencing efficiency and thereby producing apparently distinct genetic dependencies; or (b) heterochromatin in the SH region is nucleated at multiple, independent sites. In the first model, RNAi or shelterin pathways trigger heterochromatin spreading across the entire SH domain up to the SH/SU boundary. Supporting this, heterochromatin has been shown to spread efficiently over comparable distances when the entire SH region has been removed (Tashiro et al. 2017), implying that spreading can occur independently of SH sequences. Similarly, in budding yeast, functional segregation of the nucleation and spreading pathways suggests that spreading can emerge from a single telomere-proximal site (Brothers and Rine 2022). Model (b), by contrast, would require previously overlooked cryptic nucleation sites internal to SH.

We currently favor this second model for several reasons. First, silencing is maintained at telomere-proximal sites in the absence of both RNAi- and shelterin-dependent nucleation pathways. Second, Clr4 remains associated with distinct, non-coding RNA-expressing loci even in the absence of RNAi and pre-existing H3K9 methylation. This pattern resembles that observed at perincentromes loci (Khanduja et al. 2024) and at heterochromatin islands (Zofall et al. 2012; Hiriart et al. 2012), where non-coding RNAs expressed from Clr4 recruitment sites promote heterochromatin nucleation independently of RNAi through the Erh1-Mmi1 complex (EMC). Third, these Clr4 binding sites overlap with persistent H3K9me3 domains in a spreading-deficient *swi6* mutant (Suppl. Figure S6) (Khanduja et al. 2024; Kennedy et al. 2024). Together, these findings support a model in which RNA-dependent, but RNAi-independent recruitment of Clr4, potentially mediated by the EMC, enables heterochromatin nucleation within the SH domain.

### Multiple epigenetic states co-exist at the subtelomere

A hallmark of heterochromatin is epigenetic variegation, in which multiple gene expression states can be inherited in a metastable manner. One early example is Telomere Position Effect (TPE), first documented in flies (Levis et al. 1985) and later studied extensively in budding yeast (Gottschling et al. 1990). Early studies using artificial minichromosomes (Nimmo et al. 1994) demonstrated TPE in *S. pombe*, and a more recent study showed metastable maintenance of subtelomeric heterochromatin using *ade6* reporters (Kawakami et al. 2024).

Our results identify at least three distinct epigenetic behaviors within the subtelomeres: First, the region surrounding 17.5 kb is tightly silenced and rapidly recovers from perturbations. We interpret these observations as evidence that this *tlh2-*proximal site efficiently recruits heterochromatin assembly factors, thereby conferring a chromatin state that is highly resistant to perturbation. This behavior is consistent with robust nucleosome assembly, high occupancy, and low turnover (Aygun et al. 2013; Cutter DiPiazza et al. 2021; Taneja et al. 2017). Second, ‘orange’ in SCS28-ΔVI^1R^ behaves in a manner most consistent with metastable epigenetic inheritance. Silencing at this site can adopt multiple states that are stably propagated across generations, and perturbations regenerate the same clonal distribution observed during initial reporter characterization. This type of metastable behavior is somewhat reminiscent of PEV in flies (Elgin and Reuter 2013). The observation that the 7.1 kb fragment missing in this reporter is also absent in the natural Tel1R structural variant suggests that this region frequently undergoes recombination. It further implies that different epigenetic states on individual chromosomal arms may co-exist, even in strains with native SH sequences. Finally, the more distal reporters in SCS33 and SCS37 show strong silencing under steadystate conditions but lose it completely after perturbation, with no recovery even after a month or upon *clr4* reintroduction. This indicates a highly fragile epigenetic state: silencing is stably maintained once established but can hardly be re-established after disruption. A similar pattern occurs at the MAT locus upon RNAi loss (Hall et al. 2002) where silencing remains stable but fails to recover after perturbation (Greenstein et al. 2018; Jia et al. 2004; Wang and Moazed 2017). This has been attributed to the inability of Clr4 to nucleate in the absence of the RNAi machinery, even though maintenance remains intact. A comparable mechanism may operate near the 33 and 37 kb regions, where nucleation or spreading factors may act infrequently, transiently, or fail to rebind acetylated chromatin.

### Unique aspects of SH heterochromatin point to a more euchromatic and unstable chromatin environment

When comparing subtelomeres with other heterochromatin loci, several aspects of SH heterochromatin stand out. One is their dynamic behavior, which occupies an intermediate position between the stable MAT locus and the highly unstable states of RNAi-driven domains (Greenstein et al. 2018; Obersriebnig et al. 2016). At SCS28-ΔVI^1R^ and SCS28, we observe infrequent ON ⟷OFF fluctuations that appear like switches between metastable states. Their detection in even a small cell population suggests switching rates much higher than those of classic bistable systems such as *MAT*Δ*K*, which occur at 10^-4^ per generation (Grewal and Klar 1996). These findings place SH heterochromatin in an intermediate, metastable category, consistent with previous reports.

Moreover, the genetic requirements of SH-heterochromatin more closely resemble those of ectopically assembled heterochromatin in euchromatin than those of constitutive heterochromatin. Specifically, we previously showed that heterochromatin spreading in euchromatin depends on the HIRA histone chaperone and the histone variant H2A.Z-depositing Swr1 complex, while being strongly antagonized by chromatin remodelers (Greenstein et al. 2022). A similar pattern emerges for SCS28-ΔVI^1R^ and SCS37, although here remodeler subunits can either promote or oppose silencing. We interpret this as evidence of greater nucleosome instability compared to constitutive heterochromatin sites, consistent with a more euchromatin-like setting. Another parallel with ectopic spreading is sensitivity to H3K4 methylation: H3K4me3 deposited by Set1 acts as a barrier to spreading when heterochromatin boundaries are removed (Greenstein et al. 2020). At SH heterochromatin, loss of the H3K4me3 demethylase Jmj2 (Huarte et al. 2007) enhances silencing of SCS28-ΔVI^1R^ and SCS37, which suggest that failure to remove H3K4 methylation imposes a barrier to spreading. Since SH heterochromatin domains lack defined DNA-encoded boundaries (Wood et al. 2002; Cam et al. 2005), such epigenetic barriers may redirect H3K9 methylation to internal sites such as the 28 kb region, thereby enhancing silencing.

Finally, SH heterochromatin competes with other heterochromatin domains for limiting silencing factors. This is evident from our observation that RNAi mutants enhance silencing at telomere-distal sites in strains lacking the common block VI or V, indicating that factors released from RNAi-dependent heterochromatin sites elsewhere are redirected to SH heterochromatin. Further, our genetic screen uniquely identified chromatin remodelers that promote silencing at SCS28-ΔVI^1R^ and SCS37, in contrast to their usual antagonistic role at other heterochromatin regions (Greenstein et al. 2022; Sahu et al. 2024). We propose that, in their absence, silencing is strengthened at constitutive heterochromatin sites, thereby diverting limiting factors away from SH regions. In line with this model, we observed that RNAi- and shelterin-dependent silencing is concentrated at telomere-proximal regions but that silencing profiles become more uniform across SH in their absence. Such spatial buffering may stabilize silencing at internal SH sites while preventing heterochromatin from spreading into the adjacent SU domain, thereby protecting stress-responsive genes from inappropriate silencing (Tashiro et al. 2017).

SH heterochromatin, while divergent in sequence across species, has globally conserved features such as its “mosaic” and repetitive nature and lack of defined boundaries, representing an example of “convergent” chromosome evolution. In *S. pombe*, we show that SH heterochromatin harbors multiple, epigenetically distinct behaviors. These differences are not dictated by the overall presence of repressive H3K9 methylation but rather reflect variation in histone acetylation and the activity of yet-to-be-defined cryptic nucleators that operate independently of RNAi or shelterin. Notably, subtelomeres are hotspots of structural variation and genome innovation, with similar alterations reported at individual subtelomeric loci in humans. Whether similar epigenetic fragmentation of subtelomeric heterochromatin occurs in other eukaryotes remains an open question. Elucidating how these position-specific silencing behaviors influence genome stability, gene regulation, and long-term adaptation will be a key goal for future studies.

## Material and Methods

### Yeast strains and media

All *S. pombe* strains, plasmids and oligonucleotides used in this study are listed in Supplementary Tables S1, S2, and S3, respectively. Gene deletions were generated by standard genome engineering procedures using transformation with polymerase chain reaction (PCR) products and genomic integration via homologous recombination (doi.org/10.1002/yea.1142). Generated strains were validated by colony PCR. Reporter strains for monitoring silencing by flow cytometry (FC) were generated by sequentially inserting three transcriptionally encoded fluorescent reporters, as described by Greenstein et al. (2018), into the subtelomeric regions of chromosome arm IIR into the *SD4[2R+]* strain from Junko Kanoh’s laboratory (Tashiro et al. 2017) using the CRISPR/Cas9-based SpEDIT method (Torres-Garcia et al. 2020).

Superfolder GFP (SF-GFPs.p., ‘green’) driven by the ade6 promoter was integrated proximal to the tlh2 gene approximately 11 kb downstream of the telomeric repeats; ade6p-driven Kusabira orange (mKO2s.p., ‘orange’) was integrated at 17.5, 28, 33 or 37 kb from the telomeric repeats; act1p-driven E2Crimson (E2Cs.p., ‘red’) was inserted at the nearest euchromatic region ∼46.5 kb (note: subtelomeric positions are corrected for the ∼4 kb sequence missing at the end of chromosome IIR in the annotated chromosome sequence on www.pombase.org). For each of the four ‘orange’ reporter insertion sites (17.5, 28, 33, or 37 kb), two biologically independent clones (A and B) were analyzed. Additional mutations (e.g., *tas3*Δ, *taz1*Δ) were introduced into these reporter strains by homologous recombination, except for *clr4* deletions, which were introduced prior to the insertion of the ‘green’ and ‘orange’ reporters.

For reverse transcriptase combined with quantitative PCR (RT-qPCR), chromatin immunoprecipitation (ChIP)-qPCR and flow-cytometry experiments, cells were grown in rich medium (Yeast Extract Supplemented, YES) at 32°C.

### RNA extraction and cDNA synthesis

RNA extraction and cDNA synthesis were carried out as previously described (Braun et al. 2011; Holla et al. 2020; Muhammad et al. 2024). As previously described, yeast cultures (50 ml, OD_600_ = 0.4– 0.8) were harvested at 4 °C and frozen in liquid nitrogen. Cell pellets were resuspended in Trizol with zirconia beads and lysed by bead beating (4 × 30 s, 5 min rests on ice). After centrifugation, RNA was extracted twice with chloroform, precipitated using isopropanol, washed twice with 75% ethanol, air-dried, and resuspended in 60 μl RNase-free water. Concentration and purity were assessed with a NanoDrop™ spectrophotometer. A 20 μg aliquot of RNA was treated with DNaseI (Ambion) for 1 h at 37 °C, and the reaction was stopped with 6 μl of inactivation reagent. For cDNA synthesis, 5 μg of DNase-treated RNA was reverse-transcribed using oligo(dT)20 primers (50 μM) and 0.25 μl SuperScript IV (Invitrogen), following the manufacturer’s instructions. cDNA was quantified by real-time PCR using PowerTrack™ SYBR Green Mastermix (Applied Biosystems™) on a QuantStudio 5 Real-Time PCR System. Primers used for qPCR are listed in Supplementary Table S3. Transcript levels were normalized to *act1* and further processed as described in the corresponding Figure Legends.

### Chromatin immunoprecipitation (ChIP)

ChIP-qPCR was performed as described previously (Braun et al. 2011; Muhammad et al. 2024; Georgescu et al. 2020). As previously described, yeast cultures (OD_600_ = 0.6–0.8) were cross-linked with 1% formaldehyde for 10 min and quenched with 150 mM glycine. Cells were washed twice with 50 ml ice-cold PBS (phosphate-buffered saline) and cell pellets were frozen in liquid nitrogen. Cells were lysed in ice-cold lysis buffer A (50 mM HEPES-KOH pH 7.5, 140 mM NaCl, 1 mM EDTA, 1% Triton X-100, 0.1% sodium deoxycholate) containing protease inhibitors and zirconia beads. Chromatin was sheared by sonication (30 s on/off cycles, 30 min, 4 °C) and incubated overnight with antibodies against H3K9me2 (Abcam ab1220), H3K9me3 (Active Motif 39161), H3K14ac (Abcam ab52946), or H4K16ac (Active Motif 39167), followed by capture with Dynabeads Protein G. Beads were washed, de-crosslinked, treated with proteinase K, and DNA was purified using a ChIP DNA Clean and Concentrator kit (Zymo Research). Input (1:100) and IP (1:20) DNA samples were quantified by real-time PCR using PowerTrack™ SYBR Green Mastermix (Applied Biosystems™) on a QuantStudio 5 system. Primers used for qPCR are listed in (Supplementary Table S3). IP signals were normalized to input and further processed as described in the corresponding Figure Legends.

### Flow cytometry

For gene expression analysis in single cells, flow cytometry (FC) analysis was performed according to a previously described protocol (Greenstein et al. 2018). *S. pombe* cells were grown to stationary phase in YES media and then diluted to a concentration of OD600 = 0.1 in YES, followed by incubation at 32°C for 4 – 5 h prior to FC analysis. The Fortessa X-50 instrument (BD), equipped with a high-throughput sampler (HTS) module, was employed for flow cytometry analysis. Sample sizes ranged from approximately 2000 to 100,000 cells, depending on the growth conditions of the respective strain. Compensation was performed using strains expressing no fluorescent proteins (XFP) and a single-color control XFP (SF-GFPsp (‘Green’), mKO2sp (‘Orange’) or E2Csp(‘Red’)). Compensated ‘Green’ and “Orange” signals were normalized to (‘Red’) expressed from a euchromatic control locus in each single cell. Additionally, a maximum expression value for ‘Green’ and ‘Orange’ was set based on their expression in a heterochromatin-deficient control strain (*clr4Δ*), where reporters should be in an ON state. However, since the reporters are inserted at heterochromatin domains, which are prone to recombination in clr4Δ strains, there is a risk of losing these reporters. To overcome this issue, color-negative cells were excluded by setting a minimum cutoff for ‘Green’ and ‘Orange’ This threshold was determined using a control strain expressing only the ‘Red’ reporter, which represents a fully repressed state for both ‘Green’ and ‘Orange’. Specifically, the “OFF threshold” was calculated as the median fluorescence intensity of the ‘Red’-only control plus 15 times the median absolute deviation (MAD). ‘Max’ expression values for ‘Green’ and ‘Orange’ were then calculated from these color-positive cells in the no-heterochromatin control strain. Subsequently, ‘Red’ normalized ‘Green’ and ‘Orange’ signals were scaled to the corresponding ‘max’ values in our analysis strains, and the scaled, normalized signals were plotted in 2D hexbin or density plots for visualization. To estimate the fraction of fully nucleated cells, cells falling below the defined ‘Green’ and ‘Orange’ OFF thresholds were gated and isolated. The fraction of these fully repressed cells was then calculated and reported as the OFF percentage in the corresponding hexbin plots.

### Flow cytometry activated cell sorting (FACS)

Cells were grown overnight in 3 ml YES media and then diluted in the morning to OD=0.1 in YES and grown for 5–6 hours (OD=0.4). Cells from 2 mL of culture were pelleted and washed once with ice cold FACS buffer (1XPBS, 1% BSA, and 5mM EDTA). Cells were re-suspended in 500µl of ice cold FACS buffer and passed through a 35-40 μm cell-strainer. An additional 1.5 mL cold FACS Buffer was added to the strainer to wash any trapped cells into the collection tube. Sorting was performed using FACSAria2 (Becton Dickinson). Cells were considered for sorting had to pass selection for (1) Young, primarily singlet G2 cells, using size gate determined by forward (FSC) and side (SSC) scatter, as described in Greenstein et al. 2018., and (2) intact heterochromatin at 17.5 kb (‘green’^OFF^), excluding all cells above the range of an unstained control. Cells that passed this double filter were then gated into ‘orange’^OFF^, and ‘orange’^HIGH^ signal defined by the following: OFF encompassed signal overlapping that of an unstained control, and HIGH encompassed signal overlapping that of the ‘orange’ only control strain PAS52, which carries *ade6p:*mKO2 at a euchromatic locus (Greenstein et al. 2018). Approximately 8^10^6^ cells from each subpopulation were collected. Immediately after sorting cells were divided into two thirds for ChIP experiments and one third for RT-qPCR, and were subjected to the appropriate fixation treatment for either experiment. ChIP experiments were performed as previously described (Greenstein et al. 2018).

### Flow cytometry-based screen

Our curated chromatin knock-out collection comprises a subset of chromatin function mutants from the *S. pombe* BIONEER (ver3) gene deletion library. SCS reporters were crossed into this library as previously described (Greenstein et al. 2022; Muhammad et al. 2024). Knock-out strains were validated in this and our previous study (Muhammad et al. 2024) using deletion-specific barcode sequences published by Han et al. (2010) together with genomic PCR analysis. Validation results are provided in Suppl. Tables S4–S6.

Scaled and normalized fluorescence intensity values for the ‘Orange’ reporter, as described above, were stratified into four discrete expression zones (Zones 1–4) using custom-defined thresholds along the Y-axis. These thresholds included an experimentally determined “OFF threshold” (orangeOFF), 0.4 and 0.8. The orangeOFF threshold was established based on a control strain expressing only the ‘Red’ reporter, which represents a fully repressed baseline state for both ‘Green’ and ‘Orange’ reporters. The “OFF threshold” was calculated as the median fluorescence intensity of the ‘Red’-only control plus 8 times the median absolute deviation (MAD). Cells were assigned to one of four zones corresponding to increasing levels of ‘Orange’ expression: OFF (Zone 1), mid-low (Zone 2), mid-high (Zone 3), and ON (Zone 4). Samples containing fewer than 1000 cells were excluded from further analysis. The fractions of cells within each zone were calculated for each sample, and the data were visualized as stacked bar plots to represent the distribution of ‘Orange’ expression states.

Mutants were clustered based on fractions obtained from zone analysis using k-means clustering. The optimal number of clusters (k) was determined in Python, with data handling performed using NumPy v2.2.6 and pandas v2.3.0. Within-cluster sum of squares (WCSS) and silhouette scores were calculated across a range of k values with scikit-learn v1.6.1 and visualized using matplotlib v3.10.3. The KneeLocator method from kneed v0.8.5 was used to identify the optimal k according to the elbow method.

### Fission Yeast Lifespan Micro-dissector (FYLM)

FYLM experiments were performed as previously described (Greenstein et al. 2018). Fluorescence was quantified as mean intensity within a region of interest. For each fluorophore (SF-GFPsp, mKOsp, and E2Csp), an empty catch tube was used to measure autofluorescence across the entire experiment. The autofluorescence was subtracted from the fluorescence traces for each cell at each frame and the ‘Green’ and ‘Orange’ to ‘Red’ ratios were calculated from these background-subtracted values. A *clr4*Δ mutant was included in each experiment to define the ON/OFF state threshold for each fluorophore ratio. The threshold for ON was defined as mean minus three standard deviations of the maximum ratio attained by *clr4*Δ cells. In Figure 4, ratios were normalized to the mean of the maximum ratios attained by each *clr4*Δ mutant cell within each experiment.

### Heat recovery assay

Freshly plated cells were inoculated into 96-well plates containing 200 µL of YES medium per well and incubated overnight with shaking (Elmi) at either 32 °C or 38 °C (Day −1). The next morning, cultures were diluted into fresh YES medium (200 µL) and maintained at the same temperature for approximately 6 h before being analyzed by flow cytometry (Day 0). Following this measurement, cultures were diluted again into YES medium and incubated overnight at 32 °C. On Day 1, cells from the overnight cultures were re-diluted into YES medium at 32 °C, grown for 6 h, analyzed by flow cytometry, and returned to the same temperature for overnight growth. This cycle was repeated for three additional days (Days 2–4).

### Trichostatin A (TSA) recovery assay

Cells from freshly streaked plates were inoculated into 96-well plates containing 200 μL YES medium supplemented with 30 μM TSA and incubated overnight with shaking (Elmi) (Day −1). On the following day (Day 0), cultures were diluted into fresh YES medium containing 30 μM TSA and analyzed by flow cytometry. Subsequently, cells were plated onto YES agar and incubated at 32°C for 2–3 days to allow recovery. Single colonies were picked, patched onto YES plates, and incubated for an additional 2–3 days at 32°C. These isolates were maintained on YES plates for a total of 12 days. On Day 12, cultures grown overnight in 96-well plates containing 200 μL YES were diluted into fresh YES medium, incubated for approximately 6 h, and analyzed by flow cytometry. Isolates were continuously maintained on YES plates, and the same growth and measurement procedure was repeated on Days 19 and 28.

## Source code availability

Flow cytometry raw data and scripts for their analysis have been deposited on Zenodo under the accession number **doi: 10.5281/zenodo.21360361**.

## Acknowledgements

We thank members of the Braun lab, Al-Sady lab, K. Ragunathan and S. Hake for fruitful discussions during the study. We thank S. Lall (Life Science Editors) for editorial assistance and critical comments on the manuscript. Furthermore, we thank M. Wilhelm for technical assistance. Schemes in Figures were created with BioRender.com. S.B. was supported by the German Research Foundation (DFG) through the Heisenberg Programme (project ID 464293512) and by the European Union through the MSCA ITN Cell2Cell fellowship to A.M. (project ID: 860675). B.A.-S. and I.J.F. were supported by National Science Foundation grant 2113319. B.A.-S. Was additionally supported by National Institutes of Health grant R35GM141888. J.K. was supported by the Japan Society for the promotion of Science (JSPS) KAKENHI [JP23H02408].

## Author contributions

Conceived and designed the study: A.M., J.K., M.M., I.J.F., B.A.-S., and S.B. Collected the data: A.M., J.C., C.G., R.Y.J.B., J.M.S., J.S.K., and A.A.A.A. Contributed reagents: J.K. Performed the data analysis: A.M., J.C., R.Y.J.B., J.S.K., R.I.J., J.J.G., B.A.-S., and S.B. Wrote the manuscript: A.M., B.A.-S., and S.B. with input from all authors.

## Supplementary Figure Legends

**Suppl. Figure S1:**
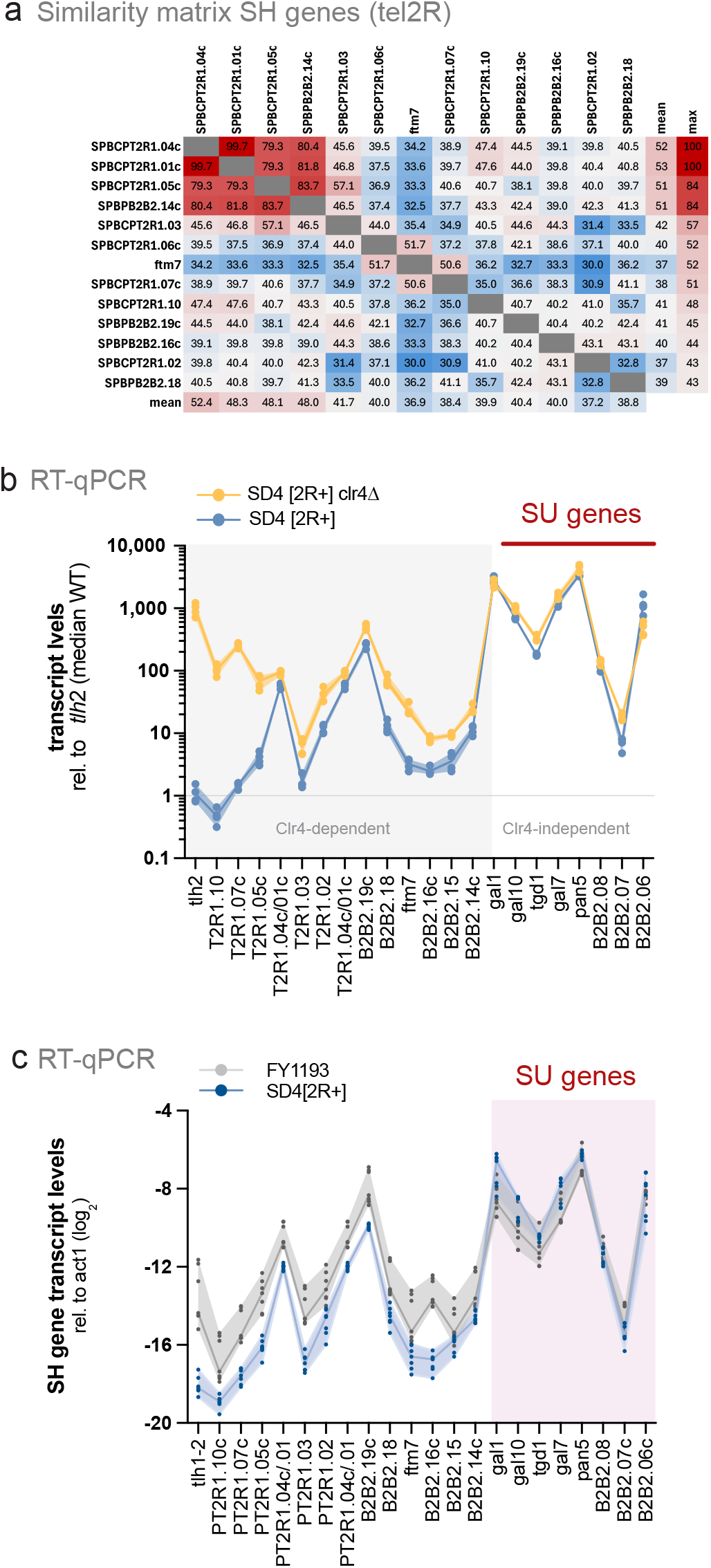
SH gene expression patterns are not impacted by number of total subtelomeres. **(a)** Similarity matrix of SH-region genes on chromosome arm Tel2R. The matrix displays pairwise nucleotide similarity scores (%) among SH genes based on sequence alignment. Rows and columns represent individual genes. The two right columns (and bottom row) display the mean and maximum similarity value, respectively, for each gene across all pairwise comparisons. **(b)** Clr4-dependence of SH and SU gene expression the *SD4[2R*^*+*^] strain. Line plot showing transcript levels by RT-qPCR of SH and SU genes in the wild-type (blue) and *clr4Δ* (yellow) *SD4[2R*+] background. Transcripts were normalized to *act1* and are shown relative to the median *tlh2*. expression level in WT samples. Independent biological replicates (*n* = 4) are shown as individual data points, with the median and range. **(c)** RT-qPCR analysis of SH and SU gene transcript levels in the *SD4[2R*+] (blue) and FY1193 (grey). FY1193 (Ekwall et al., 1999) is a derivative of the common laboratory strain 972 and retains all native subtelomeric arms. Transcript levels were quantified using strain-specific genomic DNA for normalization to account for differences in SH copy number and were subsequently normalized to *act1* (log_2_ scale). Independent biological replicates (*n* = 7) are shown as individual data points, with the median and range.

**Supplementary Figure S2:**
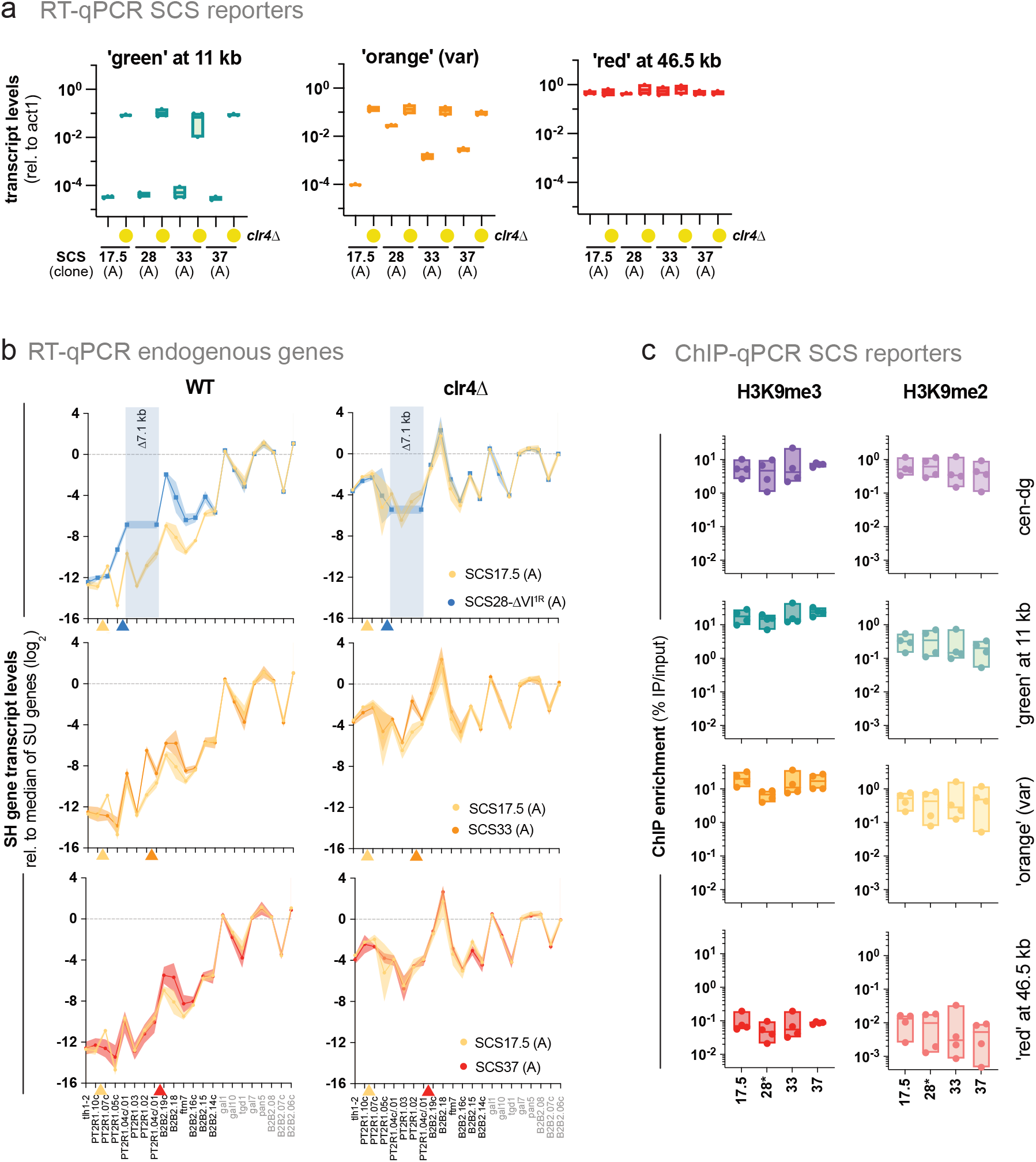
Characterization of gene silencing and genetic architecture of single-cell reporter strains. **(a)** RT–qPCR analysis of reporter transcript levels in wild-type and *clr4*Δ strains across all SCS constructs. Transcript abundance was measured for the ‘green’ reporter (11 kb), ‘orange’ reporters (17.5, 28, 33, and 37 kb), and the ‘red’ reporter (46.5 kb), and normalized to *act1*. Each data point represents an independent biological replicate (*n* = 3). Data from clone A are shown. **(b)** RT–qPCR analysis of transcript levels for genes within the SH and SU regions in SCS17.5 (yellow) compared with SCS28-DVI^1R^ (blue; top), SCS33 (orange; middle), and SCS37 (red; bottom) reporter strains (clone A) in wild-type (left panel) and *clr4*Δ (right panel) backgrounds. Transcript levels were normalized to *act1* and are presented relative to the median expression of SU-region genes (log_2_ scale). Normalization to the median SU-region expression provides a robust internal reference. Data represent independent biological replicates (WT strains: SCS17.5, *n* = 14; WT SCS28-ΔVI^1R^, *n =* 15, WT SCS33, *n = 11;* WT SCS37, *n* = 9; *clr4*Δ strains: *n* = 4) and are shown as the mean with 95% confidence interval (shaded area). The schematic (below left) displays gene positions in the SH and SU regions. The ∼7.1-kb region absent from SCS28 clone A (SCS28-DVI^1R^) is shaded in blue. **(c)** ChIP–qPCR for H3K9me3 (left) and H3K9me2 (right) enrichment, shown as IP/input without additional normalization (cf. Figure 2e). From top to bottom, panels correspond to cen *dh-dg*, the ‘green’ reporter, the ‘orange’ reporter, and the ‘red’ reporter.

**Supplementary Figure S3:**
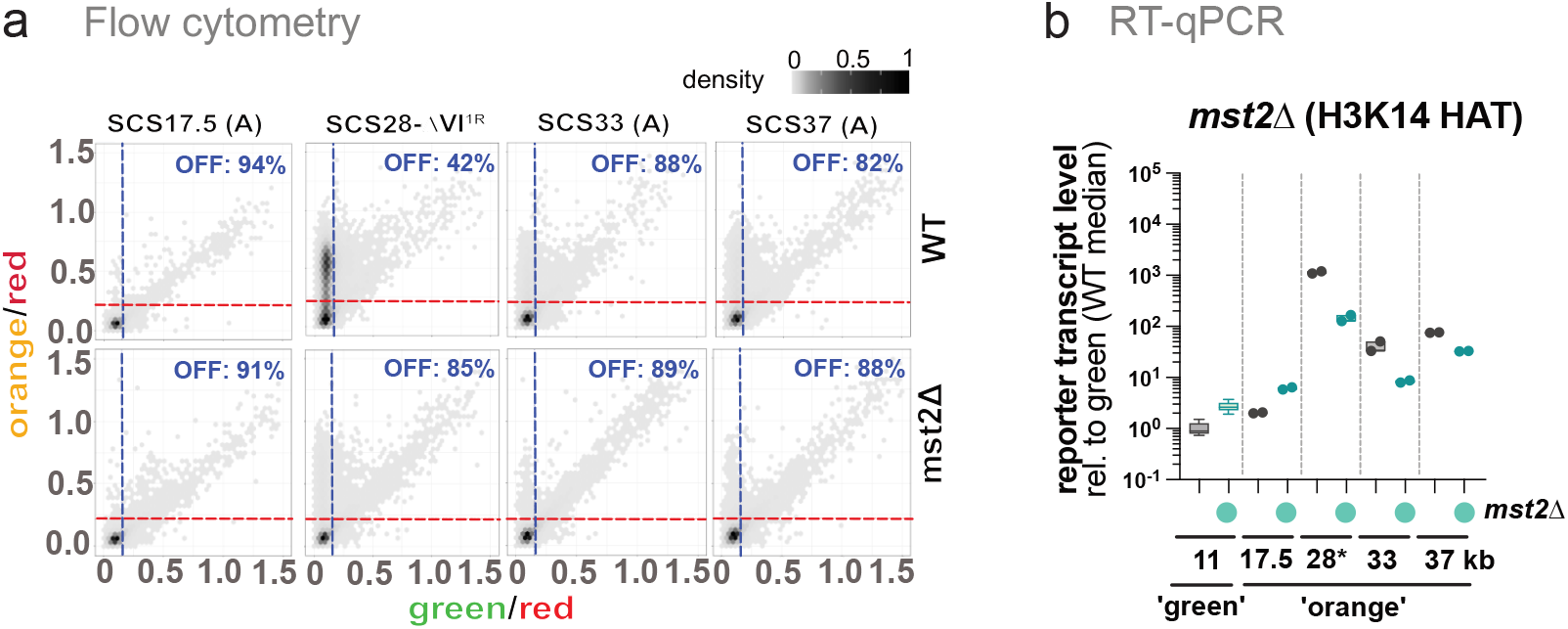
Dependence of SCS reporter silencing on Mst2. **(a)** Two-dimensional hexbin density plots showing red-normalized ‘green’ (x-axis) and ‘orange’ (y-axis) fluorescence for SCS reporters in WT (top) and *mst2*Δ cells (bottom). Blue and red dashed lines indicate OFF thresholds for green and orange reporters, respectively. **(b)** RT–qPCR analysis of reporter transcript levels in WT and *mst2*Δ cells. Transcript levels of the ‘green’ reporter (11 kb) and ‘orange’ reporters (17.5, 28-DVI^1R^, 33, and 37 kb) were measured in wild-type (grey) and *mst2Δ* (green) strains. Transcript levels were normalized to *act1* and are shown relative to the median ‘green’ reporter expression in WT samples. Data from independent biological replicates are shown as box-and-whisker plots (Tukey) for ‘green’ (*n* = 8) and as individual data points for each ‘orange’ reporter (*n* = 2).

**Supplementary Figure S4:**
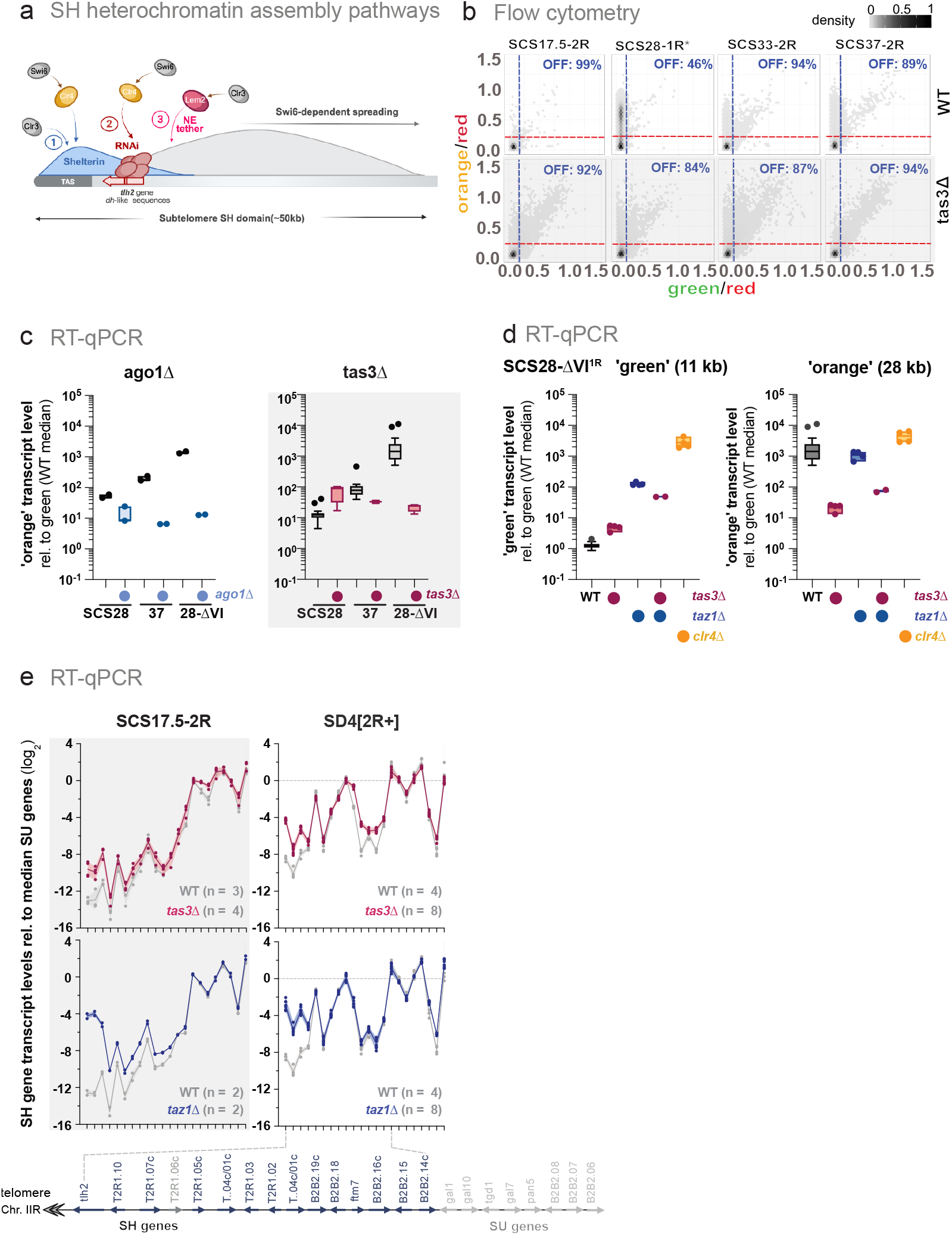
Dependence of SCS reporter silencing on RNAi and shelterin pathways. **(a)** Schematic representation of heterochromatin nucleation pathways at the telomeric and subtelomeric SH region (∼50 kb). The diagram illustrates the Shelterin-dependent pathway (blue), RNAi-mediated pathway (red), and Lem2-dependent nuclear tethering pathway (pink). Heterochromatin spreading is indicated as Swi6-dependent. Figure created with BioRender. **(b)** Two-dimensional hexbin density plots showing red-normalized green (x-axis) and orange (y-axis) fluorescence for SCS reporters in WT (top) and *tas3*Δ cells (bottom). Blue and red dashed lines indicate OFF thresholds for green and orange reporters, respectively. The shaded area shows reproduced data from Figure 2 (see Figure 2 for details). **(c)** RT–qPCR analysis of ‘orange’ transcript levels in *ago1*Δ. Left: *‘*orange’ transcript levels in wild-type or *ago1*Δ in SCS28, SCS37 and SCS28-DVI^1R^. Transcript levels were normalized to *act1* and are presented relative to the median ‘green’ reporter expression in WT samples. The shaded panel on the right reproduces corresponding *tas3*Δ versus WT from Figure 3b for comparison. Data represent two independent biological replicates. **(d)** RT–qPCR analysis of reporter transcript levels in the SCS28-DVI^1R^ strain across different genetic backgrounds. Transcript levels of the ‘green’ reporter (11 kb) and ‘orange’ reporter (28 kb) were measured in WT (grey), *tas3Δ* (red), *taz1Δ* (blue), *tas3Δ taz1Δ* (red and blue), and *clr4Δ* (yellow) strains. Transcript levels were normalized to *act1* and are presented relative to the median ‘green’ reporter expression in WT samples. Data are shown as box-and-whisker plots (Tukey) from independent biological replicates (WT, *n* = 16; *tas3*Δ, *n* = 4; *taz1*Δ, *n* = 4; *tas3*Δ *taz1*Δ, *n* = 2; *clr4*Δ, *n* = 4). Data for WT, *tas3*Δ, *taz1*Δ, and *clr4*Δ single mutants are reproduced from Figure 3b. **(e)** RT–qPCR analysis of transcript levels across the SH and SU regions in different genetic backgrounds. The line plots show gene expression in SCS17.5-2R (left) and *SD4[2R+]* (right) strains. Transcript levels were measured in wild-type (grey), *tas3Δ* (red), and *taz1Δ* (blue). Transcript levels were normalized to *act1* and are shown relative to the median expression of SU-region genes (log_2_ scale). The number of independent biological replicates is indicated in each plot. The schematic (below left panel) shows the organization of SH and SU genes. In the left panel, the light grey shading indicates data reproduced from Figure 3c.

**Supplementary Figure S5:**
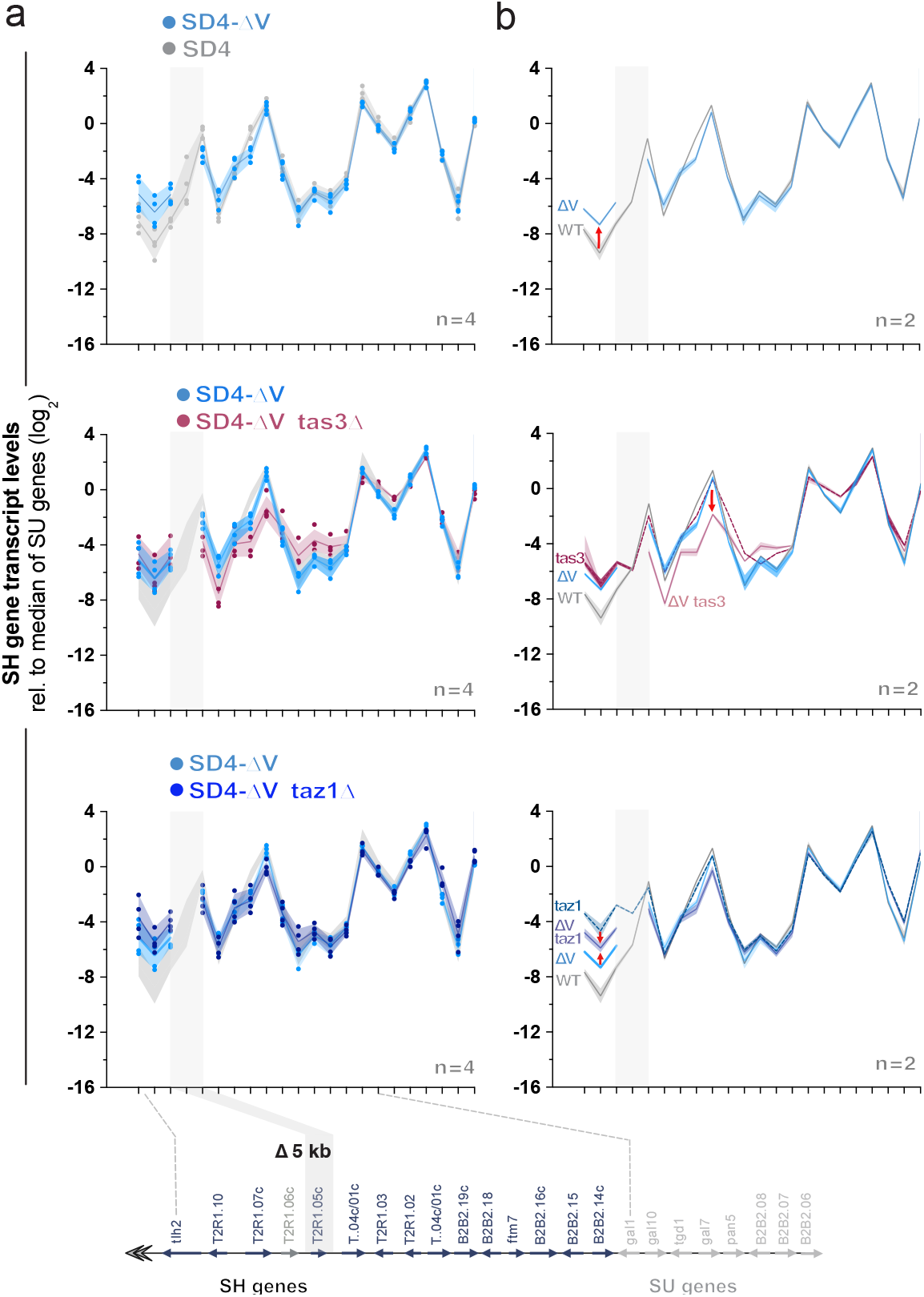
Block V locally control gene silencing at the subtelomere in the parental *SD4*[*2R*^*+*^] strain. **(a**,**b)** RT-qPCR analysis of the effect of the 5-kb block V deletion on endogenous subtelomeric gene expression in the parental *SD4[2R*^*+*^] strain lacking reporter insertions. Top: *SD4[2R*^*+*^] WT (grey) versus *SD4[2R*^*+*^] ΔV (light blue). Middle: *SD4[2R*^*+*^] ΔV (light blue) versus *SD4[2R*^*+*^] ΔV *tas3*D (dark red). Bottom: *SD4[2R*^*+*^] ΔV (light blue) versus *SD4[2R*^*+*^] ΔV *taz1*D (dark blue). Transcript levels were normalized to *act1* and are shown relative to the median expression of SU-region genes (log_2_ scale). For (a), four biological replicates from two independent experiments are shown as individual data points, together with the mean and range. For (b), two biological replicates from one independent experiment are shown as the mean and range. Data for the WT and ΔV strains are reproduced from panel (a). In addition, the middle and bottom panels in (b) include the corresponding *tas3*Δ and *taz1*Δ single mutants lacking the block V deletion (dotted lines). Red arrows indicate loci with substantial expression differences between the corresponding strains.

**Supplementary Figure S6:**
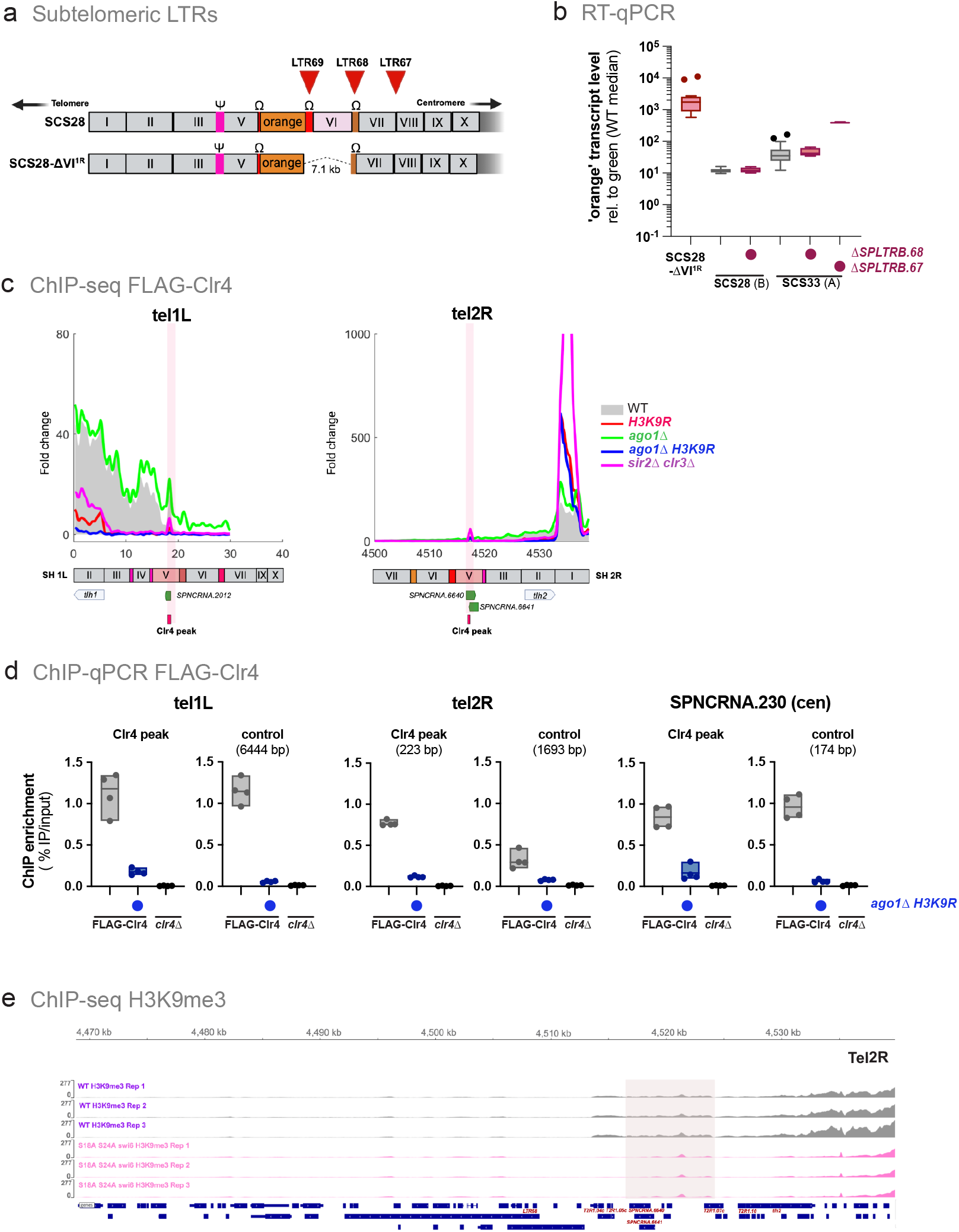
Control of SH gene silencing by *cis-*elements outside known nucleation sites. **(a)** Schematic of the location of Long Terminal Repeats (LTRs, red arrows) in SCS28 and SCS28-DVI^1R^ **(b)** RT–qPCR analysis of ‘orange’ reporter transcript levels in SCS28-DVI^1R^, SCS28 and SCS33 strains in presence and absence of SPLTRB.68 and SPLTRB.67. Transcript levels were normalized to *act1* and are shown relative to the median green reporter expression in WT (log_10_ scale). Data are displayed as Tukey box-and-whisker plots from independent biological replicates (SCS28-DVI^1R^, *n* = 16; SCS28, *n* = 10; SCS28SPLTRB.68Δ, *n* = 4; SCS33, *n* = 29; SCS33 SPLTRB.68Δ, n = 6; and SCS33 SPLTRB.67Δ, *n* = 4). Data for SCS28-DVI^1R^, SCS28, and SCS33 are identical to those in Figure 2c. **(c)** 3xFLAG-Clr4 ChIP-seq profiles at Tel1L (left) and Tel2R (right) in WT and mutant strains. Schematics of SH regions are shown below with common sequence blocks (Oizumi et al. 2021) and the positions of Clr4 peaks and ncRNAs containing potential Mmi1-recognition sites. ChIP-seq data are reproduced from Khanduja et al. 2024 (NCBI GEO submission GSE 223202, GSE 223203). The Clr4 ChIP signal is plotted after subtracting the ChIP signal of the 3xFLAG tag alone (*clr4*Δ*::3xflag*). **(d)** 3xFLAG-Clr4 ChIP-qPCR analysis in WT and the indicated mutant strains. N-terminally tagged Clr4 was expressed from its endogenous locus; the control clr4Δ strain expressed only the triple-FLAG tag. Data shown are IP/input without additional normalization from four independent biological replicates. Clr4 peaks within subtelomeric arms at *Tel1L* and *Tel2R* correspond to peak positions shown in (c); positions of the control regions are indicated (distance in bp from peak). **(e)** H3K9me3 ChIP-seq enrichment at subtelomeric regions on Tel2R in wild-type and *swi6-S18A,S24A* mutant strains. Input-normalized ChIP–seq data from Kennedy et al. 2025 (NCBI GEO submission GSE271394) are shown for four biological replicates per genotype. The *swi6-S18A,S24A* mutant exhibits a defect in heterochromatin spreading but retains nucleation at specific loci. Chromosomal coordinates follow annotation in PomBase. A light pink box marks H3K9me3 peaks retained in the mutant background near the SPBCPT2R1.07c. Normalized bigWig tracks were visualized using Integrative Genomics Viewer (IGV); grey denotes wild-type, pink denotes the *swi6-S18A,S24A* mutant.

**Supplementary Figure S7:**
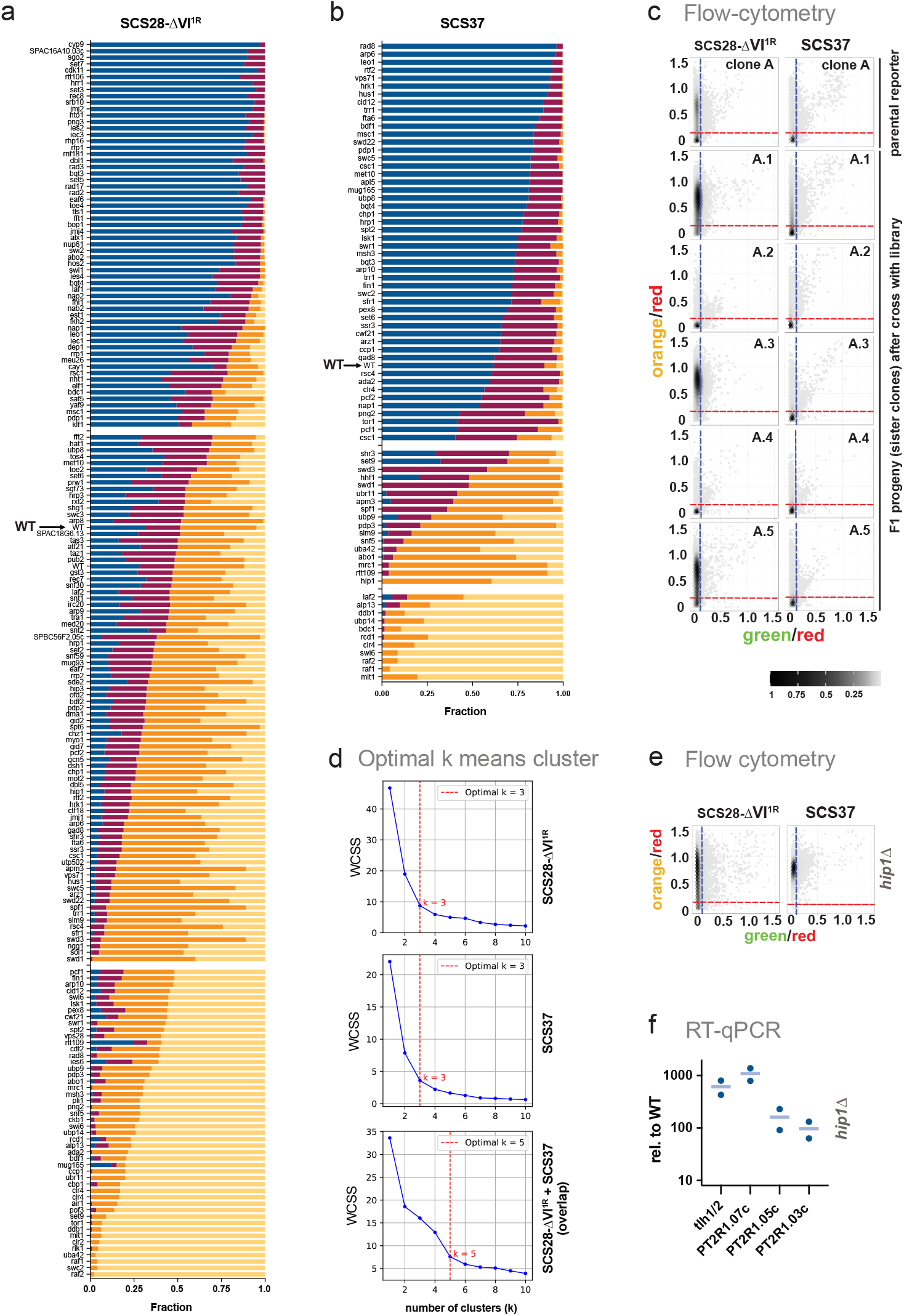
Validation of targeted fluorescence-based screen and clustering. **(a)** and **(b)** Stacked bar plots showing the distribution of cells across four fluorescence expression zones (zone 1–4, increasing orange fluorescence normalized to red) in wild-type and mutant strains in the SCS28-DVI^1R^ (panel a) or SCS37 (panel b) reporters. The genes identified the individual screens was grouped into three clusters each based on zone distribution using optimal k-means clustering (k = 3; see c). For each cluster group, horizontal bars represent individual mutants, ordered from top to bottom by increasing fraction of cells in zones 3 and 4 for each cluster group. Wild-type is indicated with an arrow. **(c)** Effect of heterogeneous subtelomeric arms generated large-scale genetic crosses of SCS reporter strains with the deletion mutant collection on reporter expression measured by flow cytometry. The top row shows the parental isolate 2D hexbin plots of SCS28-DVI^1R^ and SCS37. The panels below show representative F1 progency from crosses with selected wild-type strains from the mutant collection. The fluorescence distributions of the F1 progeny closely match those of the corresponding parental reporter strains (Figure 2) and strains recovered after TSA treatment (Figure 7), indicating that heterogeneous subtelomeric arms introduced by the crosses do not measurably affect the reporter readout. **(d)** Determination of the optimal number of clusters for k-means clustering based on expression profiles from individual SCS28-DVI^1R^, SCS37 screen, and the combined dataset. Panels show the within-cluster sum of squares (WCSS) plotted against the number of clusters (k) using the Elbow method. The optimal number of clusters is indicated by a red dashed line. The top, middle, and bottom rows correspond to SCS28-DVI^1R^, SCS37, and the merged dataset (SCS28-DVI^1R^ and SCS37), respectively. Based on the Elbow method, the optimal number of clusters is k = 3 for the individual SCS28-ΔVI^1R^ and SCS37 screens, and k = 5 for the combined dataset. **(e)** Flow cytometry analysis of *hip 1*Δ in SCS28-DVI1R and SCS37. Compare to Figure 5b. **(f)** RT-qPCR analysis of selected endogenous SH genes in WT and *hip1*Δ cells in the SCS28 (clone B) strain background. Transcript levels were normalized to *act1* and are shown relative to WT expression levels. Data represent two independent biological replicates from a single experiment.

**Supplementary Figure S8:**
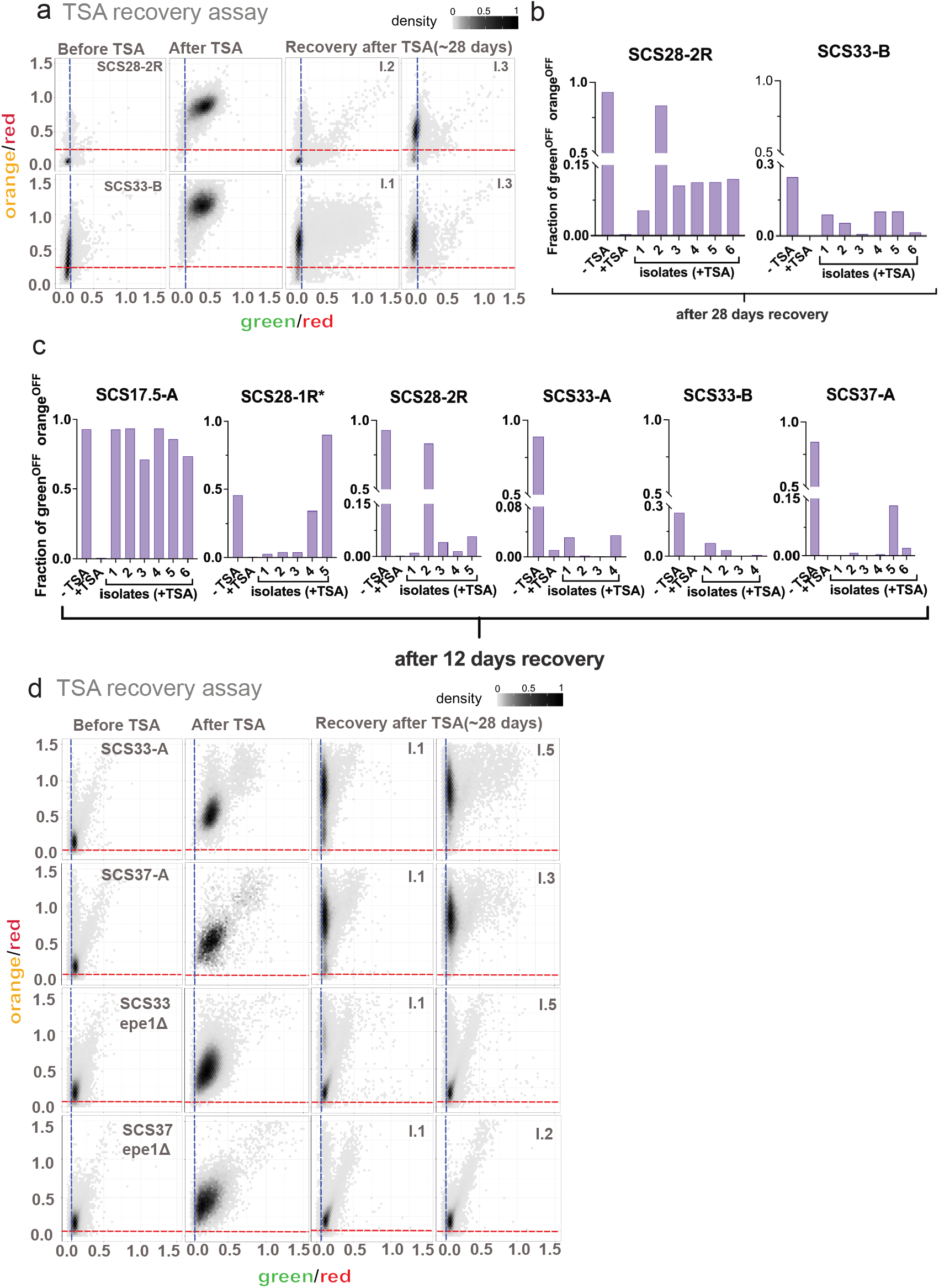
Epe1 is required for the apparent fragility of telomere-distal SH heterochromatin. **(a)** Two-dimensional hexbin density plots showing green (x-axis) and orange (y-axis) fluorescence normalized to ‘red’ for SCS reporter constructs. Blue and red dashed lines denote OFF thresholds for green and orange reporters, respectively, as in Figure 4b. Columns show fluorescence profiles in untreated wild-type (WT), following treatment with 30 μM TSA, and in two independent isolates after 28 days of recovery in nutrient-rich media. Rows correspond to SCS28 and SCS33 constructs (clone B) containing orange reporters inserted at 28 or 33 kb; green and red reporters are fixed at 11 kb and 46.5 kb, respectively. **(b, c)** Bar plots showing recovery dynamics of SCS reporter clones following TSA treatment. The y-axis represents the fraction of cells in the ‘orange’ OFF / ‘green’ OFF fluorescence state. Conditions include untreated cells, cells treated with 30 μM TSA, and isolates recovered for 28 days (b) and 12 days (c) in nutrient-rich medium in the reporter strains as indicated. For each SCS reporter, 6 independent post-recovery isolates are shown. **(d)** Two-dimensional hexbin density plots showing green (x-axis) and orange (y-axis) fluorescence normalized to ‘red’. Blue and red dashed lines mark OFF thresholds for the ‘green’ and ‘orange’ reporters, respectively, as in Figure 6b. Columns show distributions in untreated cells, after treatment with 60 μM trichostatin A (TSA), and in two independent isolates following 28 days of recovery in nutrient-rich media. Rows correspond to SCS33 and SCS37 (‘orange’ reporter inserted at 33 kb and 37 kb, respectively; clone A) and the corresponding *epe1Δ* mutant. ‘Green’ and ‘red’ reporters are located at 11 kb and 46.5 kb, respectively.

